# Occurrence of ordered and disordered structural elements in postsynaptic proteins supports optimization for interaction diversity

**DOI:** 10.1101/448100

**Authors:** Annamária Kiss-Tóth, Laszlo Dobson, Bálint Péterfia, Annamária F. Ángyán, Balázs Ligeti, Gergely Lukács, Zoltán Gáspári

**Affiliations:** Faculty of Information Technology and Bionics, Pázmány Péter Catholic University, Práter u. 50A, 1083, Budapest, Hungary; 3in-PPCU Research Group, Esztergom, 2500-Hungary

**Keywords:** Postsynaptic density, Protein-protein interaction, Intrinsically disordered proteins, Diversity of Potential Interactions

## Abstract

The human postsynaptic density is an elaborate network comprising thousands of proteins, playing a vital role in the molecular events of learning and the formation of memory. Despite our growing knowledge of specific proteins and their interactions, atomic-level details of their full three-dimensional structure and their rearrangements are mostly elusive. Advancements in structural bioinformatics enabled us to depict the characteristic features of proteins involved in different processes aiding neurotransmission. We show that postsynaptic protein-protein interactions are mediated through the delicate balance of intrinsically disordered regions and folded domains, and this duality is also imprinted in the amino acid sequence. We introduce Diversity of Potential Interactions (DPI), a structure and regulation based descriptor to assess the diversity of interactions. Our approach reveals that the postsynaptic proteome has its own characteristic features and these properties reliably discriminate them from other proteins of the human proteome. Our results suggest that postsynaptic proteins are especially susceptible to forming diverse interactions with each other, which might be key in the reorganization of the PSD in molecular processes related to learning and memory.

## 1. Introduction

Synaptic signal transduction is an elaborate process that not only provides the means of transmitting the excited state of one neuron to another but also plays an important role in basic phenomena underlying neural development, learning and memory [1,2]. The basic cellular phenomenon of memory, long-term potentiation (LTP) results in the strengthening of synaptic connections. The postsynaptic density (PSD) is a key structure in this process, as an essential part of excitatory chemical synapses. It is composed of a dense network of proteins and provides an intricate link between the intracellular parts of membrane receptors and adhesion molecules and the cytoskeleton [3]. PSD composition and organization changes during development and the individual history of the particular neuron [4,5], leading to morphological differences between PSDs in different brain regions [6]. Recent results point to a reorganization of the PSD during sleep by the exchange of Homer protein isoforms and contributing to synaptic scaling [7,8]. The emerging view is that the PSD is continuously remodeled and has a characteristic dynamics that is manifested not only by the addition and elimination of components over time but also by dynamic restructuring while retaining its composition [9–11].

From a structural point of view, interactions can be formed via a diverse range of structural elements. Protein intrinsic disorder is defined by the lack of ability of a particular protein/segment to adopt a stable three-dimensional structure. Disordered segments can play many different roles, including the mediation of protein-protein interactions (PPIs), during which they may get fully or partially folded (folding upon binding [12]) or retain a considerable degree of disorder (fuzzy complexes [13]). Disordered segments have been shown to play important roles in the postsynaptic scaffold (PSC) proteins [14], signaling pathways [15–17] and be at key positions within interaction networks [18]. The organization of synaptic proteins also exhibits a high amount of ordered domain interactions, resulting in macromolecular assemblies occupying the PSD. The highly organized assembly of PSD strongly suggests the presence of several intermolecular interacting sites with an elaborate and thoroughly regulated distribution of occupied and unavailable or available partner binding sites with a high level of redundancy. The PSD can most likely be imagined as a supramolecular association capable of integrating and transmitting signals via reorganization. The underlying mechanisms likely include competitive binding events, allostery, and cooperativity tightly regulated through post-translational modifications (PTMs) [5,19], short linear motifs [20], and alternative splicing [21].

Recently, the role of liquid-liquid phase separation (LLPS) in the organization of the PSD has been suggested [22,23]. It has been shown that appropriate combinations of selected PSD proteins result in the formation of droplets [24,25]. The presence or absence of specific proteins and/or binding sites was shown to substantially influence the phase separation properties of PSD proteins [25]. To date, LLPS is most extensively studied for RNA-binding nuclear proteins, where RNA is an important component of the supramolecular associations exhibiting phase separation. It should be noted that RNA molecules are abundant in dendrites as a pool for in situ translation [26]. A key feature of proteins capable of LLPS is multivalency, the presence of multiple partner binding sites that can be folded domains or linear motifs in intrinsically disordered regions (IDRs) [27]. Bioinformatics analysis of phase separation is rather laborious, as the first databases containing proteins or regions driving LLPS are just being developed [28], protein-protein interactions can be reliable investigated using various tools and databases, which may open prospects to the hallmark of LLPS [29,30].

Most current descriptions of human postsynaptic proteome characterize PSD proteins using experimental procedures [31] or literature-based collection extended with gene annotation services [32]. To our knowledge, there is no comprehensive computational analysis focusing on the structural organization of PSD proteins. In this paper we use a wide range of bioinformatics methods to describe the structural characteristics of PSD proteins, focusing on features that may directly contribute to the synaptic plasticity through the formation of (PPIs).

## 2. Materials and Methods

### 2.1 Datasets

The human proteome was downloaded from UniProt [33] (2018_June release). For synaptic proteins, the SynaptomeDB [34] database and its classification of protein localization were used, defining four sets: postsynaptic, presynaptic, presynaptic active zone and vesicle-associated proteins. An additional PSD-related set was defined based on a simple search on the UniProt website with the term ‘postsynaptic scaffold human’. Proteins of the immunome were extracted from the Immunome Knowledge Base [35]. The list of nuclear proteins was taken from the supplementary material of Frege et al. [36], whereas the set of histone methylases were extracted from the accompanying data of Lazar et al. [37]. Lists of interacting protein pairs were taken from the BioPlex 2.0 database [38].

### 2.2 Annotation and prediction of protein properties

Amino acids were classified into different groups, based on their physicochemical properties: hydrophobic (A, I, L, M, V), aromatic (F, W, Y), polar (N, Q, S, T), positively charged (H, K, R,), negatively charged (D, E), rigid (P), flexible (G), and covalently interacting (C). AAIndex was used to obtain various amino acid scales [39]. To reduce the number of features, the ward.d2 function of R was used for K-means clustering [40], then based on the elbow method 5 clusters were selected to represent the features (Supplementary Figure 1). The following amino acid scales were selected randomly, each from one of the clusters: NAKH900104, KHAG800101, JUNJ780101, HUTJ700103, FAUJ880103 (for explanation see Supplementary Table 1).

Disorder prediction was made in a way that aims to minimize false positive hits arising from the inclusion of oligomeric fibrillar motifs [41]. First, the consensus of two disorder prediction methods, IUPred [42] and VSL2B [43] were determined. In the second step, all residues predicted to be in oligomeric fibrillar motifs were eliminated from the set of disordered residues [41]. Oligomeric fibrillar motifs were determined using a permissive prediction, namely, residues to form coiled coils as predicted either by COILS [44] or Paircoil2 [45], single *α*-helices as identified using FT_CHARGE [46] or collagen triple helix as obtained by HMMER [47] with the Pfam HMM for collagen (ID: PF01391.13) segments in Pfam [48]. The remaining set of residues is considered to be a good representation of segments being disordered under cellular conditions.

Binding regions within disordered segments were predicted with Anchor [42], low complexity regions with SEG [49], transmembrane (TM) regions with CCTOP [50,51], and signal peptides were predicted using SignalP [52]. Linear motifs participating in binding events were retrieved from the ELM database [53], considering only “LIG/DOC” classes. The ELM prediction was also utilized, considering only those hits that passed the various filters as described by Gouw et al. [53]. Protein domains were taken from UniProt (2018_June release), PTMs were downloaded from PhosphoSitePlus [54] and were also predicted by NetPhos [55]. Venn diagrams were created using Venny [56].

All calculated values for each protein are available in Supplementary Table 1.

### 2.3 Statistical analysis

We calculated the mean and standard deviation of these features on all datasets listed in 2.1. Furthermore, to compare the statistics of the proteome to Synaptome, PSD and PSC we also used exclusive sets. Due to the unbalanced distribution of proteins in various datasets we used bootstrapping and random sampled all datasets 1000 times. The size of these samples was derived from the smallest dataset: 80% of protein belonging to PSC was used as reference. The calculations were used to show the enrichment of features in different datasets compared to the proteome, by applying logarithmic scale on their proportion.. Means and variances for different groups are available in Supplementary Table 2. To further confirm the significance of the differences in the distributions of features, we also performed Kolmogorov-Smirnov tests for all sets vs. the full human proteome (i.e. we compared the distribution of features between the proteome and the different subsets). This data is available in Supplementary Table 3. We also counted the number of PSD and non-PSD proteins with given features above and below their respective mean values (calculated for the proteome) and performed X-square tests (with Yates correction) on the contingency tables resulted from this procedure (Supplementary Table 4).

To investigate the co-occurrence of ordered and disordered structural elements in proteins of the PSD, we counted proteins with different individual structural entities and their combinations. Significance of this data is confirmed by randomly selecting 80% of data 1000 times (as described above) and calculating the mean and standard deviations. Whenever the mean ± (1,2 and 3 fold) standard deviations did not overlap, we accepted the result to be significant with p<0.32, p<0.05 and p<0.01, respectively (Supplementary Table 5).

Similar analysis was used to analyze the possible connection between PPIs and PTMs (Supplementary Table 6).

For Diversity of Potential Interactions (DPI, see 3.5) and PPI values of the different subsets Spearman’s rank correlation was calculated.

### 2.4. Machine learning

As a first step we preprocessed the data used for machine learning by removing homologous sequences and assigning labels. For redundancy filtering CD-HIT [57] was used in an incremental manner, filtering identical proteins to 90, 70, 50 and finally to 40% identity. The remaining sequences were used to train the predictor. Labels were assigned based on the SynaptomeDB annotation.

For machine learning a feed forward neural network was developed, using one hidden layer with 40 neurons and stochastic gradient descent. Due to the non-proportional data for training and testing, bootstrap aggregating (BAGGING) was used. In each step, ten down-sampled sets were created and used to calibrate the Artificial Neural Networks (ANNs). For the final prediction, the results of individual ANNs were aggregated and weighted based on their reliability (defined as the output neuron probability). Benchmarking was done using ten-fold cross-validation and independent datasets.

Two different predictors were built: the first uses all annotations presented in Supplementary Table 1. The second predictor only uses features that can be derived from the amino acid sequence and do not depend on the annotation of different databases (shown with a grey background in Supplementary Table 1). Training and testing data are available in Supplementary Table 7.

## 3. Results

### 3.1 Datasets

In this paper, we investigated three nested subsets of proteins to the human proteome (21766 proteins): all proteins from the synaptome (1891 proteins), proteins localizing into the postsynaptic density (1761 proteins), and postsynaptic scaffold proteins (51 proteins). Three additional control datasets were defined: histone methylases (52 proteins), proteins from the nucleus (180 proteins) and the immunome (834 proteins).

According to our definition (derived from SynaptomeDB), there is considerable overlap between the synaptic proteome and proteins of the PSD, and their characteristics are on par in every case. Therefore, conclusions drawn for PSD are valid for the synaptome too, even if it is not explicitly declared.

### 3.2 Sequences feature a mix of disorder and order promoting properties

General sequence properties often help to gain insight about structural features of proteins. As a very first step we compared the length distribution of proteins in the PSD to other proteins of the proteome: on average proteins in the PSD are longer (the average length of proteins is 524 and 705 residue in the proteome and in the PSD, respectively) (Figure 1, Supplementary Figure 2). We also calculated the average amino acid content of proteins. On the one hand, these values seem similar and their variances are high, therefore these properties cannot be used alone to reliably distinguish proteins in different localizations. On the other hand, these differences are mostly significant and according to the distribution of structural domains (see 3.3) it is clear that in some level, the function and structure of proteins are imprinted in their amino acid sequence: PSD proteins have some preference to include more charged residues (∼28% and ∼25%, in the PSD and in the proteome, respectively) and to avoid cysteines (∼2% and ∼3%, in the PSD and in the proteome, respectively). These properties alone may promote disorder content, however, on some level, the lower proline content (∼5% and ∼6%, in the PSD and in the proteome, respectively) suggest globular domains may also emerge. Low complexity regions (LCRs) often, but not always, coincide with intrinsically disordered segments: interestingly proteins in PSD are not enriched in such regions (see Kolmogorov-Smirnov test: Supplementary Table 3; X-square test: Supplementary Table 4). These results may hint a balanced presence of intrinsically disordered regions (IDRs) and ordered globular domains in the PSD, however to unambiguously prove our hypothesis specialized predictions and analysis were performed (see 3.3).

**Figure 1.**
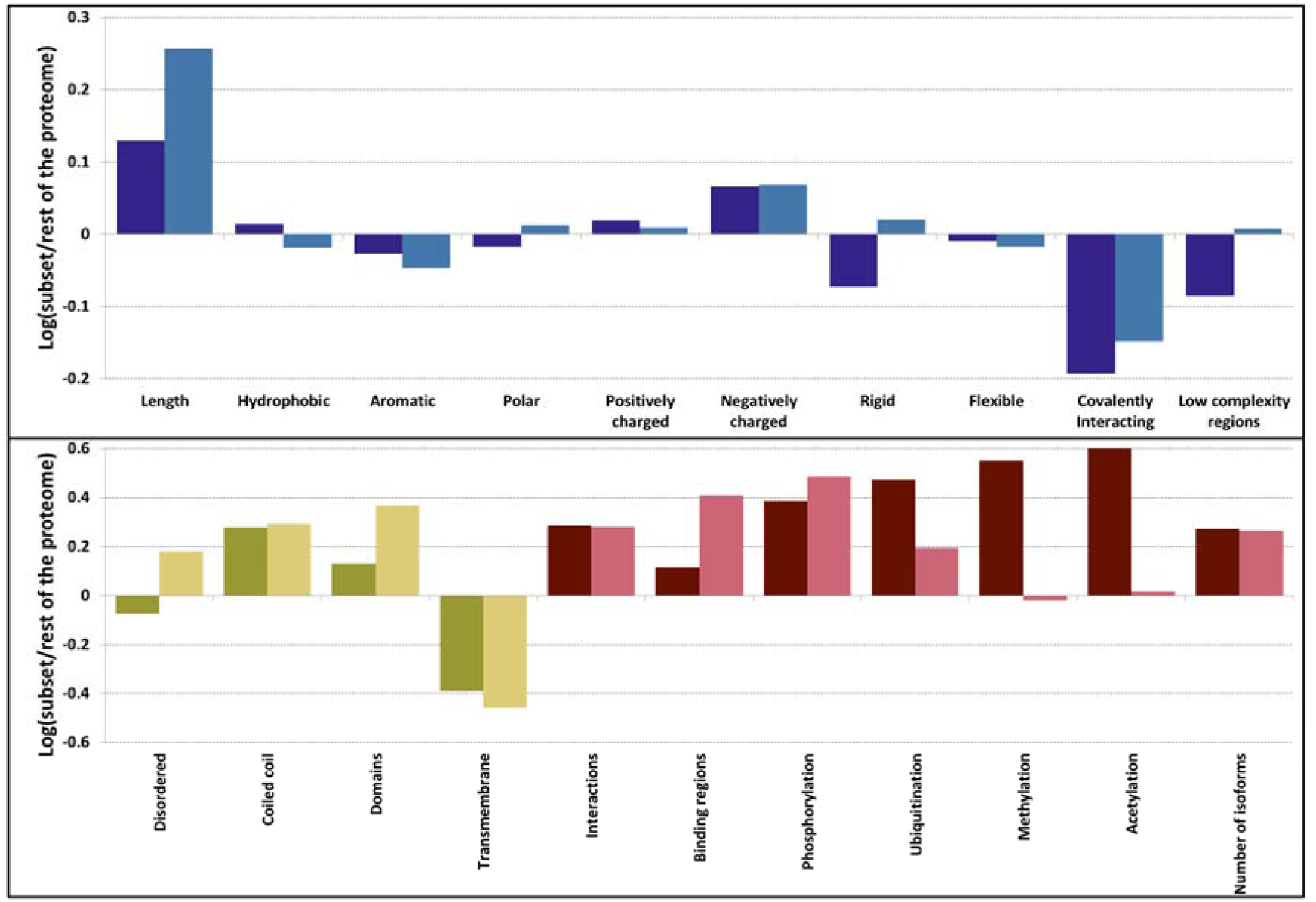
Difference sequence (blue), structure (yellow) and function (red) related properties in PSD (darker shade) and PSC (lighter shade) compared to the human proteome. Amino acids were grouped as hydrophobic (A, I, L, M, V), aromatic (F, W, Y), polar (N, Q, S, T), positively charged (H, K, R), negatively charged (D, E), rigid (P), flexible (G), and covalently interacting (C).

PSC proteins are a subset of PSD proteins with somewhat distinct features. We observed their average length is even higher compared to proteins from PSD, and almost the double of proteins of the proteome (on average 970 residue length). In contrast to the PSD, they contain a lot of IDRs Other characteristics of PSC proteins are on par with PSD proteins.

### 3.3 PSD proteins exhibit a diverse range of structured elements

As sequence data indicate differences compared to the proteome, various prediction methods were utilized to reveal different types of structural elements. We classified segments as disordered regions and ordered globular domains; then we extended this classification to groups often falling outside the classical “globular-disordered” partition: coiled-coils, a structural element already described in various PSD proteins and transmembrane segments assumed to play important role in transmitting information outside the cell. On the one hand, PSD proteins operate with a similar amount of disorder content as other proteins from the proteome (Figure 1). On the other hand, PSD proteins contain a large number of coiled coils and structured domains. We observed that not only the number of structured domains is higher, but flexible linker regions are also generally shorter in proteins of the PSD, allowing more dense placements of domains along the polypeptide chain (Supplementary Figure 3). Note that our stringent structure prediction pipeline (see Methods) has an emphasis on discriminating disordered and coiled-coil regions. Therefore the commonly occurring cross predictions between them are expected to only moderately bias these results. Another distinctive feature is the distribution of transmembrane proteins: proteins composed of seven transmembrane helices are greatly missing from the PSD (Supplementary Figure 4).

PSC proteins are on par with PSD proteins according to most of these features, however, on a more extreme way, utilizing even more coiled coils and domains. As expected from sequence characteristics, they utilize a larger amount of IDRs (Figure 1).

Another notable trend is the mutual presence of ordered and disordered segments in PSD proteins (Table 1). While the PSD exclusive proteome outnumbers PSD proteins in single-domain proteins (i.e. containing a single coiled-coil, intrinsically disordered segment, a single domain or embedded in the membrane) or in proteins where multiple IDRs or other elements appear independently, in the PSD these elements seem to emerge in a more coordinated fashion, likely to promote more diverse possibilities for PPI formation (also see significance test in Supplementary Table 5).

**Table 1.** Occurrence of structural elements in proteins (* marks significant co-occurrences, see Supplementary Table 5). IDR: Intrinsically Disordered Region; CC: Coiled-coil, TM: Transmembrane; DOM: Domain).

|  | Number of proteins |  | Proportion |  |
| --- | --- | --- | --- | --- |
|  | Proteome | PSD | Proteome | PSD |
| IDR* | 2851 | 128 | 0.14 | 0.07 |
| CC* | 1187 | 231 | 0.06 | 0.13 |
| TM* | 3460 | 154 | 0.17 | 0.08 |
| DOMAIN* | 2614 | 187 | 0.13 | 0.11 |
| IDR+CC* | 1194 | 132 | 0.06 | 0.07 |
| IDR+TM | 1224 | 92 | 0.06 | 0.05 |
| IDR+DOMAIN* | 2288 | 221 | 0.11 | 0.13 |
| IDR+CC+DOMAIN* | 1088 | 229 | 0.05 | 0.13 |
| ALL* | 125 | 21 | 0.01 | 0.01 |
| All other combination of ordered domains* (CC, TM and Domain) | 3884 | 366 | 0.19 | 0.21 |

### 3.4 The overlap between protein-protein interactions and post-translational modifications hint a tightly regulated protein network in PSD

PSD proteins operate with a high amount of PPIs (Figure 1, Supplementary Figure 5), compared to the human proteome. Interestingly, proteins from the PSD are not particularly enriched in IDRs, a commonly used element to mediate PPIs. This is also reflected in the relatively low number of predicted disordered binding regions. The extremely high amount of phosphorylation and ubiquitination sites suggest that post-translational regulation may play an essential role in PSD proteins.

As the increased number of IDRs and binding regions suggest, PSC proteins may use different mechanisms to promote PPIs. In the case of phosphorylation and ubiquitination, the trends are similar to PSD proteins. However, methylation and acetylation values are more close to the proteome used as a reference.

Since we do not have information about the specific binding regions and the interacting partners, we tried to characterize the nature of these interactions by observing their co-occurrence with proteins containing IDRs and PTMs (i.e. they emerge in the same protein, but they do not necessarily overlap). We assessed the mutual presence of PPIs, PTMs and protein disorder in the proteins of the PSD. Protein disorder for PPI formation may be somewhat less frequently employed mechanism in the PSD compared to the human proteome (i.e., ∼49% and ∼53% of interacting proteins do not contain any IDR at all in the proteome and the PSD, respectively). However, PTMs may regulate the emerging PPIs more tightly (i.e., only ∼98% compared to ∼92% of proteins participating in interactions have phosphorylation or ubiquitination site in the PSD and the proteome, respectively) (Figure 2). All these differences between the proteome and the PSD are significant according to mean and standard variation values (Supplementary Table 6).

**Figure 2.**
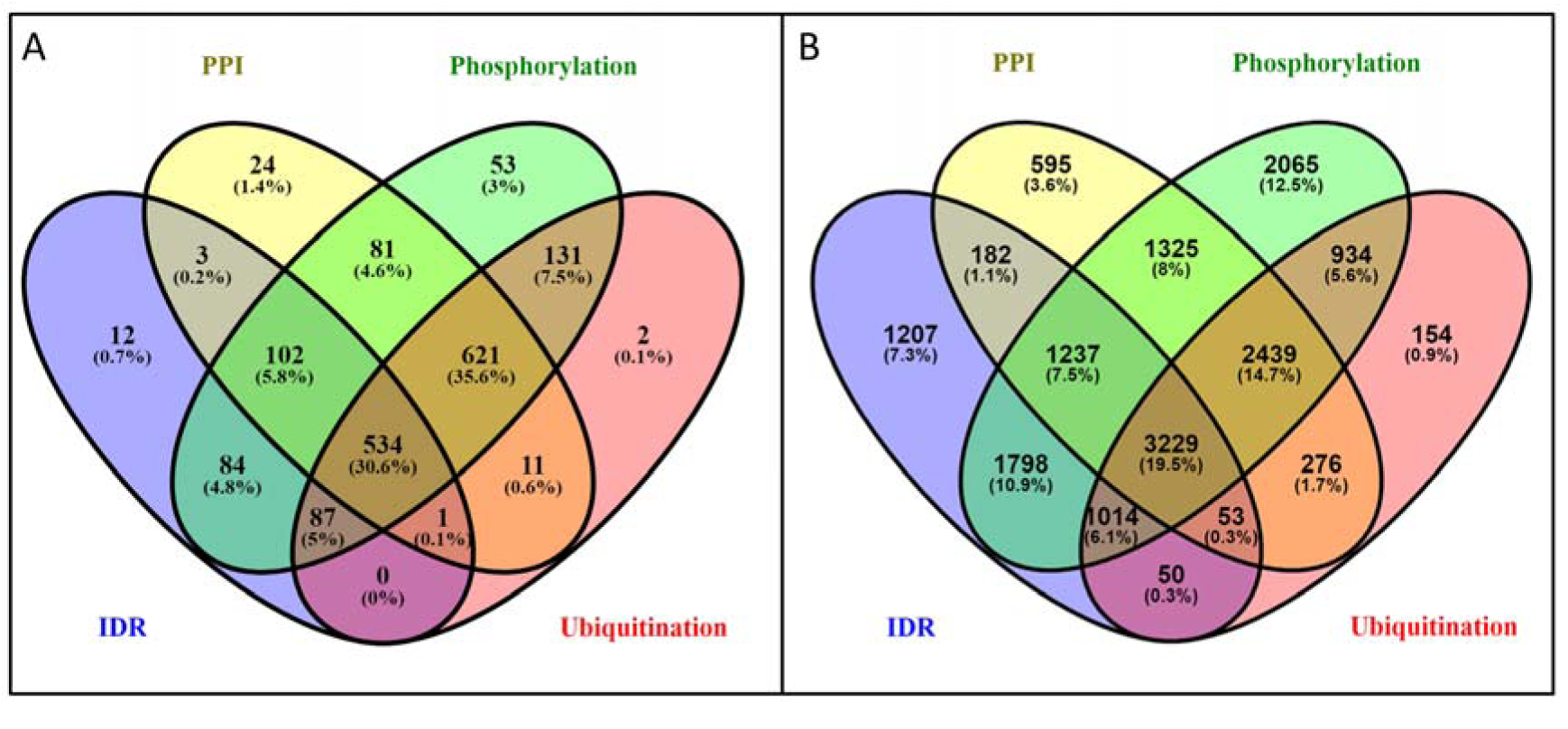
Venn diagram of proteins utilizing IDRs and PTMs to establish PPIs. A) PSD; B) human proteome. All the differences are significant with P<0.05.

Alternative splicing can also contribute to regulation, by inclusion or exclusion of binding regions and rewiring PPI networks. PSD and PSC proteins both seem to utilize this mechanism with an increased amount of isoforms compared to the human proteome (Figure 1).

### 3.5 Potential of the proteins to be engaged in multivalent interactions

Proteins in the PSD are capable of establishing a high amount of interactions, which heavily depends on the number of basic building blocks: coiled-coils can multimerize to build protein complexes, while IDRs, transmembrane proteins, and globular domains also can form inter- or intramolecular interactions. To estimate the bias caused by overlapping definitions, we calculated the overlap between these elements: we found that less than 1% of them overlap on residue level. We defined Diversity of Potential Interaction (DPI) as a composite descriptor and introduced the following elements in the equation:

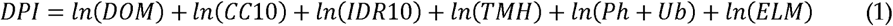

where DOM is the number of domains, CC10 is the number of coiled-coil regions longer than ten residues, IDR10 is the number of IDRs longer than ten residues, and TMH is the number of transmembrane segments, Ph and Ub are the number of phosphorylations and ubiquitination sites along the protein sequence and ELM is the number of short linear motifs aiding binding processes (LIG/DOC classes of the ELM database). These features and the calculated DPI have somewhat better distinctive properties than residue distribution (Supplementary Figures 6-13) and applying Spearman’s rank correlation indicates a significant relationship between PPIs and DPI (S. correlation=0.35; p-value<0.01). Using DPI as measure PSD proteins are on par with PSC proteins, as expected from the number of interactions (see Figure 1 and Figure 3), while the proteome has rather lower values in both cases.

**Figure 3.**
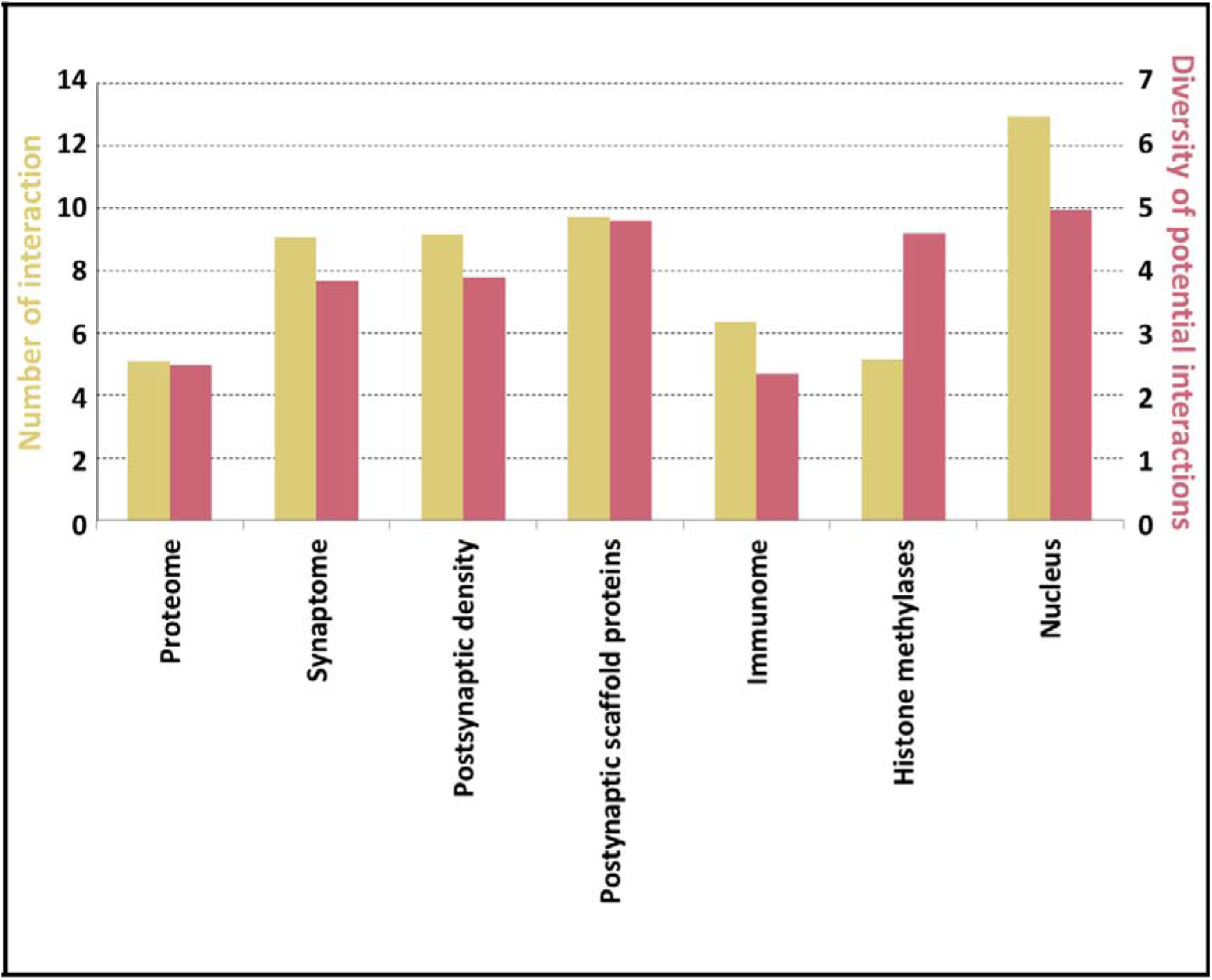
Average DPI (red) and number protein interaction values (yellow) in different protein sets.

To have a better picture how these descriptors behave, we also included three control sets with distinct features: immunome as a set of primarily globular multidomain proteins, histone methylases, known to have high IDR content and a more comprehensive list of proteins from the nucleus, shown to operate with a high amount of PPIs. We calculated the Spearman’s rank correlation between the descriptors and the average number of interactions in the selected protein sets. DPI fairly correctly estimates the potential to the formation of complexes (correlation: 0.71; p-value<0.1).

### 3.6 Sequential and structural features discriminate PSD proteins from other proteins

Although some of the above-discussed trends indicate that PSD proteins have distinctive properties, to demonstrate the prediction power of these features we established an artificial neural network (ANN). The input of the ANN consists of the calculated features discussed above. The output is a binary classification defining the localization of the protein (i.e., localizing in the PSD/not). The distribution of the positive and negative labels was not proportional; therefore we used bootstrap aggregation on the data. Briefly, this means that random samples of the data are used to train several predictors, and the final result is provided by aggregating the various outputs. The approach also helps to deal with the relatively noisy data, as in some cases it is expected that the same proteins localize inside and outside the PSD too. The algorithm achieved moderately high accuracy by incorporating all of the features presented in previous chapters (Area Under Curve (AUC): 0.84) (Figure 4 and Table 2).

**Table 2.**
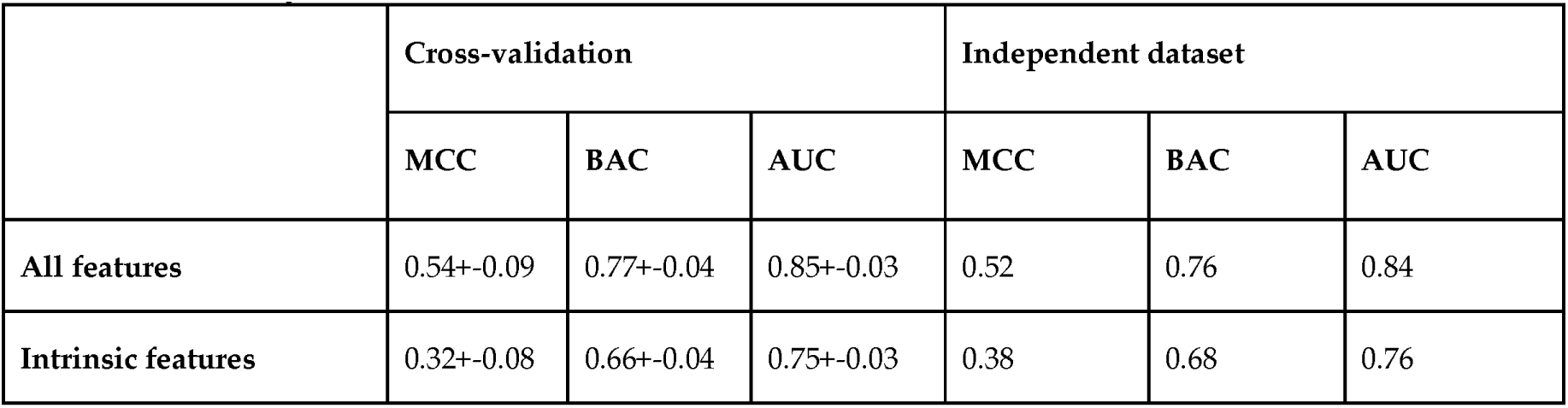
Prediction accuracy of the Neural Network. (MCC: Matthew Correlation Coefficient, BAC: Balanced Accuracy, AUC: Area Under Curve)

|  | Cross-validation |  |  | Independent dataset |  |  |
| --- | --- | --- | --- | --- | --- | --- |
|  | MCC | BAC | AUC | MCC | BAC | AUC |
| All features | 0.54+-0.09 | 0.77+-0.04 | 0.85+-0.03 | 0.52 | 0.76 | 0.84 |
| Intrinsic features | 0.32+-0.08 | 0.66+-0.04 | 0.75+-0.03 | 0.38 | 0.68 | 0.76 |

**Figure 4.**
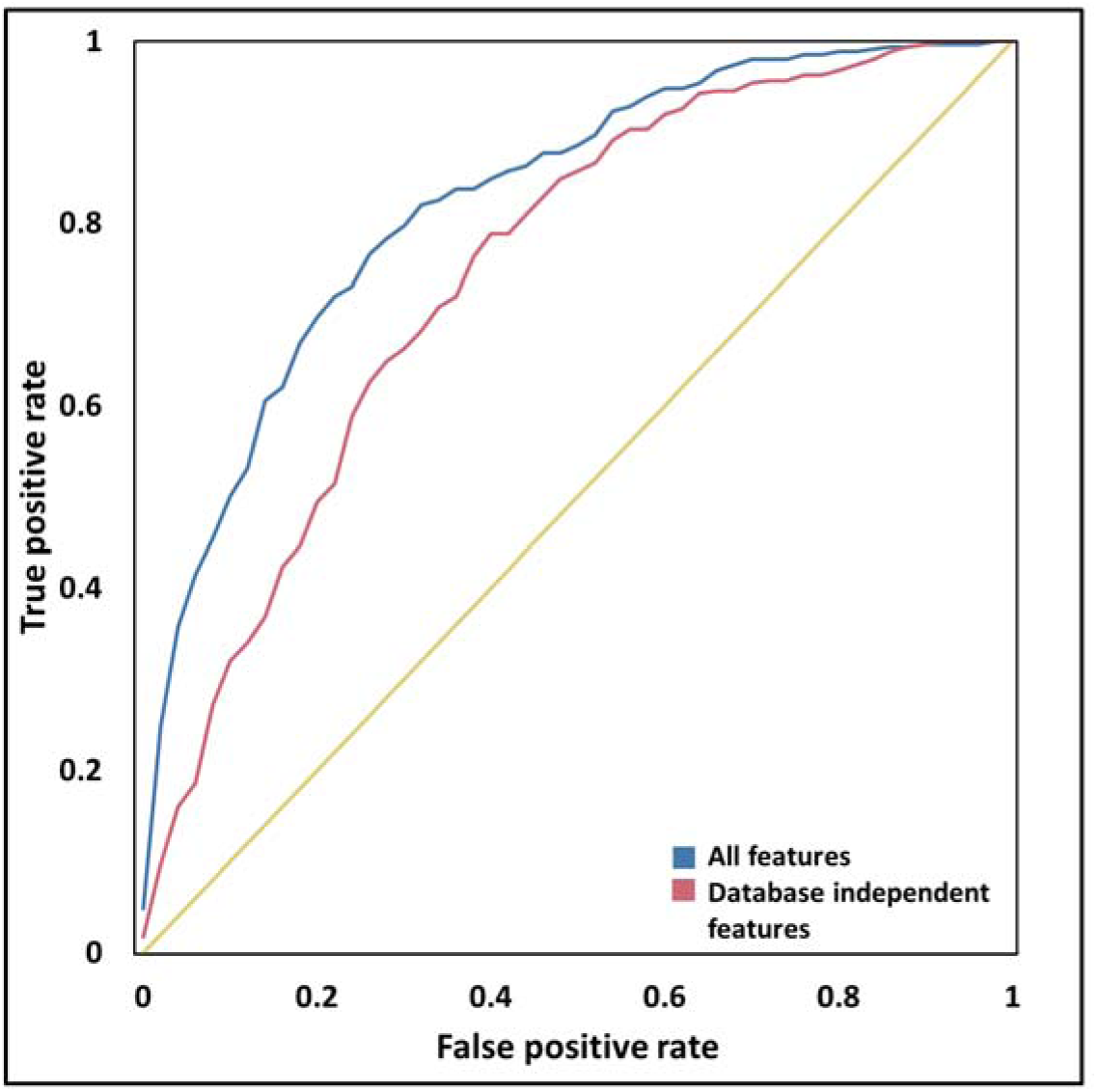
Receiver operating characteristics of ANN predictors.

Some of the characteristics used as input are database dependent, reflecting a bias in our knowledge; therefore it is hard to assess how the ANN works on proteins with more/less annotation. To overcome this problem, a second predictor was built, in which the feature space was reduced to contain only those characteristics that can be estimated based on the amino acid sequence of the proteins (intrinsic features). In some cases, annotations were replaced by prediction (e.g., phosphorylation). Although the performance of the prediction decreased, these features alone still reliably discriminate PSD proteins from others (AUC: 0.76).

Besides AUC we also calculated Matthew Correlation Coefficient and Balanced Accuracy. We observed similar values during the cross-validation and on the independent dataset. The results suggest the selected structural features are descriptive for PSD proteins and the prediction method is quite robust regarding both the features and sample sets.

## 4. Discussion

Synaptic plasticity is facilitated by proteins of the PSD, most plausibly by using their functional repertoire provided by the ability to reorganize PPIs dynamically. Our results indicate that PSD and especially PSC proteins have an increased potential to form diverse interactions and their features can be used to discriminate PSD proteins from the rest of the proteome.

General sequence properties are often used to characterize particular protein subsets, as they open prospects to structural features. Grouping amino acids based on physicochemical properties highlights exciting trends. The increased number of charged residues would hint a higher extent of intrinsic disorder; however, the low amount of prolines seems to counterbalance this effect. PSC proteins have similar characteristics, however with higher proline content that may promote intrinsic disorder.

In a recent study protein interactions were analyzed based on the structural state of participating partners, revealing major differences between “classical” protein disorder, and complexes formed by various partners. Three basic types of complexes were distinguished: autonomous folding and independent binding (i.e., the binding of two or more ordered proteins), coupled folding and binding (where an ordered protein stabilize an IDP partner) and mutual synergistic folding (interactions formed exclusively by disordered proteins) [58]. According to sequence analysis, the amino acid content of the constitutive partners in the different groups have their own characteristic, and they also differ from those described in “classical” disordered protein papers. Besides sequence analysis, differences were also shown in the bound structure: distinct groups exhibit different secondary structures, unique residue/atomic level interaction features and energy properties. They also have different biological roles in different subcellular localizations. The overall amino acid content of PSC proteins shows high correlation with the group composed of proteins going through coupled binding and folding. In contrast, proteins from PSD are more close to the “mutual synergistic folding” (MSF) class (Supplementary Figure 14). We also noted that the predicted IDR content of PSD proteins lags behind that of the proteome (Figure 1). These observations raise the question whether PSD proteins utilize a different flavor of intrinsic disorder to establish interactions. A possible scenario might be that such flexible regions are overlooked by disorder prediction methods, as they lacked MSF protein sets for training. We note that coiled coils can be regarded as a subclass of MSF complexes, however our pipeline explicitly detects them and they do not show enrichment compared to PSC proteins (Figure 1). Thus, it is plausible that MSF mechanisms other than coiled-coil formation also contribute to complex formation in the PSD.

The structural organization of the PSD utilizes a balanced distribution of different types of elements promoting PPIs. Coiled-coils, domain-domain interactions, intrinsically disordered regions, and transmembrane proteins all contribute to the formation of protein complexes in the PSD. Notably, in PSD proteins, IDRs are often paired with with other structural elements, in contrast to the full proteome, where this association is much less frequent. Although the common presence of TM segments and IDRs shows an exception (also see [59]), other combinations between IDRs and ordered segments are significantly higher. By exhibiting a diverse range of structural segments as binding regions PSD proteins likely involved in a wide range of different types of interactions to maintain the PPI network of the PSD.

Besides statistics presented in the results, detailed structure-function studies provide experimental evidence on how PPIs are formed and maintained in the PSD. Below we discuss different aspects of PSD organization by using specific well-characterized proteins as demonstrative examples.

The Shaker channel is a voltage-dependent potassium channel responsible for conducting depolarizing potassium currents when the membrane potential increases [60]. The C-terminal tail of the channel is in random coil state and contains a PDZ domain recognition motif. The motif assists the interaction with the PSD-95 scaffolding protein, helping the channel to cluster at unique membrane sites, which is important for the proper assembly and functioning of the synapse [61]. Anchor [62] can relatively accurately predict such interactions, also described as coupled binding and folding [63]. Both amino acid content (Supplementary Figure 14) and Anchor predictions indicate that PSC proteins are enriched in disordered binding regions relative to the proteome, while the number of unique partners lags behind this enrichment (Figure 1/b). The stacking of WW domain and MAPK binding motifs was described earlier in the case of TANC protein [64], also supporting the idea that PSC proteins “allow” their partners to pick from multiple unoccupied sites to fine-tune their function. In contrast, this enrichment is less pronounced in PSD and does not follow the enrichment in the number of interacting partners. These results theorize PSD proteins either more heavily utilize competitive binding with alternative partners binding to the same region, or promote different interaction modes. Since our former suggestion is rather hard to assess using computational methods, we collected other possible interaction modes between proteins.

Activity-regulated cytoskeleton-associated protein (Arc) is a crucial regulator of long-term synaptic plasticity. The protein is highly modular and contains a flexible C-terminal tail [65]. Arc oligomerization is aided by the terminal IDRs, leading to the assembly of a structure reminiscent to an HIV capsid. This way Arc can encapsulate RNA and can mediate their transfer, which was shown to play a role in cell to cell communication in the nervous system [66]. Coiled-coils are also important oligomerization motifs, linking two or more partners with high specificity and a wide range of stability [67]. The PSD protein Homer can form high-order complexes with a diverse range of partners. A tetrameric form resulting in a coiled-coil develops in a two-step process: first, the C-terminal ∼70 residues of the proteins form a parallel dimer, then two Homer dimers serve as a base for the antiparallel tetramer structure [68]. This arrangement is long enough to connect elements through the thickness of the PSD, serving direct connection between the plasma membrane and intracellular proteins through EVH1 domains binding to various scaffolding partners [69]. The mechanisms mentioned above share the characteristic feature of forming complexes by natively unfolded regions.

One can argue whether computational tools cross predict IDRs and coiled-coils, leading to a bias in our observations. To overcome this problem, our pipeline uses coiled coils as a filter when predicting IDRs. However, the lack of abundance of IDRs in PSD proteins could be explained by the fact, that we classified them as coiled coils. Calculating the overlap between IDR and coiled-coil containing proteins, and proteins forming interactions confirmed that this is not the case (Supplementary Figure 15), IDR and coiled-coil containing proteins do not dominate PPIs in PSD proteins.

PPIs depend on structural elements providing binding sites, however, they are also intensively regulated through the spatiotemporal control in the cell, often using post-translational modifications to precisely modulate the properties of the protein.

PSD-95 is a scaffolding protein in the synaptome, and the Serine at 561 position was shown to be subject to phosphorylation and work as a molecular switch. Phosphomimetic (Ser -> Asp) mutation promotes an intramolecular interaction between the guanylate kinase (GK) and the SH3 domains, inducing a highly dynamic, yet closed conformation with buried binding sites. In contrast, mutating Ser561 to Alanine leads to a stable open conformation, facilitating the interaction between PSD-95 and its partners [70]. Besides such simple mechanism phosphorylation is often used cumulatively to produce an electrostatic effect [71] to fine-tune PPIs [72] or in a combinatorial way, where specific patterns can be responsible for regulating synaptic activity [73].

Additional regulation phenomena may provide further functional diversity. Using alternative splicing, different isoforms displaying characteristic sets and distributions of interaction sites may carry out distinct functions, as it was described in the case of PDZ-containing proteins [21]. Short Linear Motifs also play a role by mediating protein-protein interactions to contribute to a plethora of functions. Their short length and structural flexibility provide plasticity to emerge by convergent evolution and fine-tune interaction networks [74].

The multivalent nature of PSD proteins can also lead to additional structure-related phenomena: the SynGAP postsynaptic protein forms a coiled-coil trimer, shortly followed by a C-terminal binding motif interacting with the third PDZ domain of PSD-95. After complex formation, the assembly goes through liquid-liquid phase separation. The enrichment of SynGap and PSD-95 was shown to highly correlate with synaptic activity [75], advocating the critical role of the formation of membraneless organelles in synaptic processes [23].

Due to the high variance of presented features, they cannot be used alone to describe PSD proteins. However, we assumed that using their combination may reveal certain aspects of PSD proteins. For this purpose we introduced descriptors to assess the presence and distribution of different elements in PSD proteins and their possible role in interactions. Since PPI formation may occur through many distinct structural elements, we included several of these to thoroughly catch many possible aspects of protein complex formation. Using Diversity of Potential Interactions (DPI), including structural and post-translational regulation members, the potential of a protein to form distinct interactions can be adequately defined. Verification with different control sets confirmed that it can be used directly on proteins sets to estimate their tendency to establish elaborate protein networks.

Our protein-level descriptions are all focused on PPIs, and thus it is not trivial whether they are discriminative features of PSD proteins, or they are only relevant in the context of specific protein complexes. Machine learning is a commonly used tool to discriminate protein subsets when the number of features and their variance is too high to overlook. Our results demonstrate the prediction power of the presented sequential, structural and regulation features. Considering the noise present in the datasets (as some proteins may be localized in the PSD and other locations too, moreover the possible false classification of source databases also adds a bias), the prediction is remarkably accurate. Applying structure based machine learning algorithm may enhance synaptome database development by expediting data collection and reducing manual effort.

In conclusion, we suggest that postsynaptic proteins and in particular, postsynaptic scaffold proteins are capable of forming diverse kinds of interactions with their partners that we propose to play a key role in the functional organization of the postsynaptic density and its dynamic rearrangements upon stimuli. We also found this ability is imprinted in the amino acid sequence and can be used to discriminate proteins with propensity to form a high number of interactions, or using machine learning to distinguish the PSD proteome from other proteins.

## Supplementary Materials

The following are available online at www.mdpi.com/xxx/s1

SMaterial: Includes Supplementary Discussion and Supplementary Figures.

SFigure 1: K-means evaluation of various numbers of clusters of AAIndex

SFigure 2: Protein length distribution (blue: PSD, red: human proteome)

SFigure 3: Flexible linker length distribution compared to the human proteome (blue: synaptome, yellow: PSD, red: PSC)

SFigure 4: Transmembrane helix distribution (blue: PSD, red: human proteome)

SFigure 5: Distribution of the number of interacting partners (blue: PSD, red: human proteome)

SFigure 6: Distribution of protein lengths in all datasets.

SFigure 7: Distribution of intrinsically disordered residue content in all datasets.

SFigure 8: Distribution of coiled-coils residue content in all datasets.

SFigure 9: Distribution of transmembrane residue content in all datasets.

SFigure 10: Distribution of the number of globular domains in all datasets.

SFigure 11: Distribution of the number of phosphorylation sites in all datasets.

SFigure 12: Distribution of the number of ubiquitination sites in all datasets.

SFigure 13: Distribution of the DPI values in all datasets.

SFigure 14: A: Correlation of amino acid content between interaction classes based on the structural state of participating partners and postsynaptic proteins. Columns: Autonomous folding and independent binding (i.e., the binding of two or more ordered proteins), coupled folding and binding (where an ordered protein stabilize an IDP partner) and mutual synergistic folding (interactions formed exclusively by disordered proteins), No folding, no binding (i.e., the “classical” disordered definition). Rows: Postsynaptic Scaffold proteins, and proteins from the postsynaptic density. Color scales from red (negative correlation) to green (positive correlation). B: Change of amino acid content of the group mutual synergistic folding (blue) and PSD proteins (red) compared to the proteome. C: Change of amino acid content of the group coupled binding and folding (blue) and PSC proteins (red) compared to the proteome.

SFigure 15: Overlap of coiled coils, IDRs and PPIs in PSD proteins.

Stable 1: Calculated properties and labels of all proteins used in the study.

Stable 2: Mean and standard deviation of calculated properties in different groups. Top: sets derived from other sources (i.e. SynaptomeDB, Uniprot etc). Middle: Exclusive human proteome datasets. Bottom: Values calculated by downsampling and bootstrapping default sets 1000 times

Stable 3: P-values of Kolmogorov Smirnov tests calculated between the proteome and different sets on all features

Stable 4: Contingency tables and calculated p-values of chi-square tests. The independent observations are: I) PSD/nonPSD II) feature is above/below of the mean calculated on the proteome

Stable 5: Co-occurence of structral formations in PSD/nonPSD proteins. Means and standard deviations were calculated for selected combinations

Stable 6: Co-occurence of PPI promoting features in the PSD and proteome proteins. Means and standard deviations were calculated for all combinations

Stable 7: List of proteins used to train and test the Artificial Neural Network.

## Author Contributions

Conceptualization, Z.G. and L.D.; Methodology, Z.G. and L.D.; Formal Analysis, A.K-T, A.A, L.D and B.P.; Investigation, A.K-T, L.D, G.L and B.L.; Data Curation, A.A. and L.D.; Writing – Original Draft Preparation, Z.G and L.D.; Writing – Review & Editing, all authors.; Visualization, L.D. and A.K-T,; Supervision, Z.G.; Project Administration, Z.G.; Funding Acquisition, Z.G.

## Acknowledgments

The constructive remarks of Bálint Mészáros and the technical assistance of Zsófia Kálmán are gratefully acknowledged.

## References

1. Rudy, J.. The Neurobiology of Learning and Memory; Sinauer Associates, Sunderland, MA, USA, 2008;

2. Ho, V.M.; Lee, J.-A.; Martin, K.C. The Cell Biology of Synaptic Plasticity. Science (80-.). 2011, 334, 623–628.

3. Feng, W.; Zhang, M. Organization and dynamics of PDZ-domain-related supramodules in the postsynaptic density. Nat. Rev. Neurosci. 2009, 10, 87–99.

4. Dosemeci, A.; Tao-Cheng, J.H.; Vinade, L.; Winters, C.A.; Pozzo-Miller, L.; Reese, T.S. Glutamate-induced transient modification of the postsynaptic density. Proc. Natl. Acad. Sci. U. S. A. 2001, 98, 10428–32.

5. Ehlers, M.D. Activity level controls postsynaptic composition and signaling via the ubiquitin-proteasome system. Nat. Neurosci. 2003, 6, 231–242.

6. Farley, M.M.; Swulius, M.T.; Waxham, M.N. Electron tomographic structure and protein composition of isolated rat cerebellar, hippocampal and cortical postsynaptic densities. Neuroscience 2015, 304, 286–301.

7. Diering, G.H.; Nirujogi, R.S.; Roth, R.H.; Worley, P.F.; Pandey, A.; Huganir, R.L. Homer1a drives homeostatic scaling-down of excitatory synapses during sleep. Science (80-.). 2017, 355, 511–515.

8. de Vivo, L.; Bellesi, M.; Marshall, W.; Bushong, E.A.; Ellisman, M.H.; Tononi, G.; Cirelli, C. Ultrastructural evidence for synaptic scaling across the wake/sleep cycle. Science 2017, 355, 507–510.

9. Blanpied, T.A.; Kerr, J.M.; Ehlers, M.D. Structural plasticity with preserved topology in the postsynaptic protein network. Proc. Natl. Acad. Sci. 2008, 105, 12587–12592.

10. MacGillavry, H.D.; Song, Y.; Raghavachari, S.; Blanpied, T.A. Nanoscale Scaffolding Domains within the Postsynaptic Density Concentrate Synaptic AMPA Receptors. Neuron 2013, 78, 615–622.

11. Meyer, D.; Bonhoeffer, T.; Scheuss, V. Balance and Stability of Synaptic Structures during Synaptic Plasticity. Neuron 2014, 82, 430–443.

12. Sugase, K.; Dyson, H.J.; Wright, P.E. Mechanism of coupled folding and binding of an intrinsically disordered protein. Nature 2007, 447, 1021–1025.

13. Tompa, P.; Fuxreiter, M. Fuzzy complexes: polymorphism and structural disorder in protein-protein interactions. Trends Biochem. Sci. 2008, 33, 2–8.

14. Tantos, A.; Han, K.-H.; Tompa, P. Intrinsic disorder in cell signaling and gene transcription. Mol. Cell. Endocrinol. 2012, 348, 457–465.

15. Dunker, A.K.; Cortese, M.S.; Romero, P.; Iakoucheva, L.M.; Uversky, V.N. Flexible nets. The roles of intrinsic disorder in protein interaction networks. FEBS J. 2005, 272, 5129–5148.

16. Vucetic, S.; Xie, H.; Iakoucheva, L.M.; Oldfield, C.J.; Dunker, A.K.; Obradovic, Z.; Uversky, V.N. Functional Anthology of Intrinsic Disorder. 2. Cellular Components, Domains, Technical Terms, Developmental Processes, and Coding Sequence Diversities Correlated with Long Disordered Regions. J. Proteome Res. 2007, 6, 1899–1916.

17. Csizmok, V.; Follis, A.V.; Kriwacki, R.W.; Forman-Kay, J.D. Dynamic Protein Interaction Networks and New Structural Paradigms in Signaling. Chem. Rev. 2016, 116, 6424–6462.

18. Cortese, M.S.; Uversky, V.N.; Dunker, A.K. Intrinsic disorder in scaffold proteins: getting more from less. Prog. Biophys. Mol. Biol. 2008, 98, 85–106.

19. Schuman, E.M.; Dynes, J.L.; Steward, O. Synaptic Regulation of Translation of Dendritic mRNAs. J. Neurosci. 2006, 26, 7143–7146.

20. Songyang, Z.; Fanning, A.S.; Fu, C.; Xu, J.; Marfatia, S.M.; Chishti, A.H.; Crompton, A.; Chan, A.C.; Anderson, J.M.; Cantley, L.C. Recognition of unique carboxyl-terminal motifs by distinct PDZ domains. Science 1997, 275, 73–7.

21. Sierralta, J.; Mendoza, C. PDZ-containing proteins: alternative splicing as a source of functional diversity. Brain Res. Rev. 2004, 47, 105–115.

22. Feng, Z.; Zeng, M.; Chen, X.; Zhang, M. Neuronal Synapses: Microscale Signal Processing Machineries Formed by Phase Separation? Biochemistry 2018, 57, 2530–2539.

23. Feng, Z.; Chen, X.; Zeng, M.; Zhang, M. Phase separation as a mechanism for assembling dynamic postsynaptic density signalling complexes. Curr. Opin. Neurobiol. 2019, 57, 1–8.

24. Zeng, M.; Shang, Y.; Araki, Y.; Guo, T.; Huganir, R.L.; Zhang, M. Phase Transition in Postsynaptic Densities Underlies Formation of Synaptic Complexes and Synaptic Plasticity. Cell 2016, 166, 1163–1175.e12.

25. Zeng, M.; Chen, X.; Guan, D.; Xu, J.; Wu, H.; Tong, P.; Zhang, M. Reconstituted Postsynaptic Density as a Molecular Platform for Understanding Synapse Formation and Plasticity. Cell 2018, 174, 1172–1187.e16.

26. Van Driesche, S.J.; Martin, K.C. New frontiers in RNA transport and local translation in neurons. Dev. Neurobiol. 2018, 78, 331–339.

27. Boeynaems, S.; Alberti, S.; Fawzi, N.L.; Mittag, T.; Polymenidou, M.; Rousseau, F.; Schymkowitz, J.; Shorter, J.; Wolozin, B.; Van Den Bosch, L.; et al. Protein Phase Separation: A New Phase in Cell Biology. Trends Cell Biol. 2018, 28, 420–435.

28. Mészáros, B.; Erdős, G.; Szabó, B.; Schád, É.; Tantos, Á.; Rawan, A.; Tamás, H.; Murvai, N.; Kovács, O.P.; Kovács, M.; et al. PhaSePro: the database of proteins driving liquid-liquid phase separation. Nucleic Acids Res. 2020, submitted.

29. Martin, E.W.; Mittag, T. Relationship of Sequence and Phase Separation in Protein Low-Complexity Regions. Biochemistry 2018, 57, 2478–2487.

30. Pritišanac, I.; Vernon, R.M.; Moses, A.M.; Forman Kay, J.D.; Pritišanac, I.; Vernon, R.M.; Moses, A.M.; Forman Kay, J.D. Entropy and Information within Intrinsically Disordered Protein Regions. Entropy 2019, 21, 662.

31. Bayés, À.; Collins, M.O.; Croning, M.D.R.; van de Lagemaat, L.N.; Choudhary, J.S.; Grant, S.G.N. Comparative Study of Human and Mouse Postsynaptic Proteomes Finds High Compositional Conservation and Abundance Differences for Key Synaptic Proteins. PLoS One 2012, 7, e46683.

32. Pirooznia, M.; Wang, T.; Avramopoulos, D.; Valle, D.; Thomas, G.; Huganir, R.L.; Goes, F.S.; Potash, J.B.; Zandi, P.P. SynaptomeDB: an ontology-based knowledgebase for synaptic genes. Bioinformatics 2012, 28, 897–9.

33. UniProt Consortium, T. UniProt: the universal protein knowledgebase. Nucleic Acids Res. 2018, 46, 2699–2699.

34. Pirooznia, M.; Wang, T.; Avramopoulos, D.; Valle, D.; Thomas, G.; Huganir, R.L.; Goes, F.S.; Potash, J.B.; Zandi, P.P. SynaptomeDB: an ontology-based knowledgebase for synaptic genes. Bioinformatics 2012, 28, 897–9.

35. Ortutay, C.; Vihinen, M. Immunome knowledge base (IKB): an integrated service for immunome research. BMC Immunol. 2009, 10, 3.

36. Frege, T.; Uversky, V.N. Intrinsically disordered proteins in the nucleus of human cells. Biochem. Biophys. Reports 2015, 1, 33–51.

37. Lazar, T.; Schad, E.; Szabo, B.; Horvath, T.; Meszaros, A.; Tompa, P.; Tantos, A. Intrinsic protein disorder in histone lysine methylation. Biol. Direct 2016, 11, 30.

38. Huttlin, E.L.; Bruckner, R.J.; Paulo, J.A.; Cannon, J.R.; Ting, L.; Baltier, K.; Colby, G.; Gebreab, F.; Gygi, M.P.; Parzen, H.; et al. Architecture of the human interactome defines protein communities and disease networks. Nature 2017, 545, 505–509.

39. Kawashima, S.; Kanehisa, M. AAindex: amino acid index database. Nucleic Acids Res. 2000, 28, 374.

40. Ward, J.H. Hierarchical Grouping to Optimize an Objective Function. J. Am. Stat. Assoc. 1963, 58, 236–244.

41. Gáspári, Z. Is Five Percent Too Small? Analysis of the Overlaps between Disorder, Coiled Coil and Collagen Predictions in Complete Proteomes. Proteomes 2014, 2, 72–83.

42. Mészáros, B.; Erdős, G.; Dosztányi, Z. IUPred2A: context-dependent prediction of protein disorder as a function of redox state and protein binding. Nucleic Acids Res. 2018, 46, W329–W337.

43. Obradovic, Z.; Peng, K.; Vucetic, S.; Radivojac, P.; Dunker, A.K. Exploiting heterogeneous sequence properties improves prediction of protein disorder. Proteins Struct. Funct. Bioinforma. 2005, 61, 176–182.

44. Lupas, A.; Van Dyke, M.; Stock, J. Predicting coiled coils from protein sequences. Science (80-.). 1991, 252, 1162–1164.

45. Berger, B.; Wilson, D.B.; Wolf, E.; Tonchev, T.; Milla, M.; Kim, P.S. Predicting coiled coils by use of pairwise residue correlations. Proc. Natl. Acad. Sci. U. S. A. 1995, 92, 8259–63.

46. Kovács, Á.; Dudola, D.; Nyitray, L.; Tóth, G.; Nagy, Z.; Gáspári, Z. Detection of single alpha-helices in large protein sequence sets using hardware acceleration. J. Struct. Biol. 2018, 204, 109–116.

47. Finn, R.D.; Clements, J.; Eddy, S.R. HMMER web server: interactive sequence similarity searching. Nucleic Acids Res. 2011.

48. Finn, R.D.; Coggill, P.; Eberhardt, R.Y.; Eddy, S.R.; Mistry, J.; Mitchell, A.L.; Potter, S.C.; Punta, M.; Qureshi, M.; Sangrador-Vegas, A.; et al. The Pfam protein families database: towards a more sustainable future. Nucleic Acids Res. 2016, 44, D279–D285.

49. Wootton, J.C. Non-globular domains in protein sequences: automated segmentation using complexity measures. Comput. Chem. 1994, 18, 269–85.

50. Dobson, L.; Reményi, I.; Tusnády, G.E. The Human Transmembrane Proteome. Biol. Direct 2015, submitted.

51. Dobson, L.; Reményi, I.; Tusnády, G.E. CCTOP: a Consensus Constrained TOPology prediction web server. Nucleic Acids Res. 2015.

52. Almagro Armenteros, J.J.; Tsirigos, K.D.; Sønderby, C.K.; Petersen, T.N.; Winther, O.; Brunak, S.; von Heijne, G.; Nielsen, H. SignalP 5.0 improves signal peptide predictions using deep neural networks. Nat. Biotechnol. 2019, 37, 420–423.

53. Gouw, M.; Sámano-Sánchez, H.; Van Roey, K.; Diella, F.; Gibson, T.J.; Dinkel, H. Exploring Short Linear Motifs Using the ELM Database and Tools. In Current Protocols in Bioinformatics; John Wiley & Sons, Inc.: Hoboken, NJ, USA, 2017; Vol. 58, p. 8.22.1-8.22.35.

54. Hornbeck, P. V.; Zhang, B.; Murray, B.; Kornhauser, J.M.; Latham, V.; Skrzypek, E. PhosphoSitePlus, 2014: mutations, PTMs and recalibrations. Nucleic Acids Res. 2015, 43, D512–D520.

55. Blom, N.; Sicheritz-Pontén, T.; Gupta, R.; Gammeltoft, S.; Brunak, S. Prediction of post-translational glycosylation and phosphorylation of proteins from the amino acid sequence. Proteomics 2004, 4, 1633–1649.

56. Oliveros, J.C. VENNY. An interactive tool for comparing lists with Venn Diagrams.

57. Fu, L.; Niu, B.; Zhu, Z.; Wu, S.; Li, W. CD-HIT: accelerated for clustering the next-generation sequencing data. Bioinformatics 2012, 28, 3150–3152.

58. Mészáros, B.; Dobson, L.; Fichó, E.; Tusnády, G.E.; Dosztányi, Z.; Simon, I. How folding and binding intertwine during protein complex formation provides an additional layer of functional regulation. J. Mol. Biol. 2019, in press.

59. Tusnády, G.E.; Dobson, L.; Tompa, P. Disordered regions in transmembrane proteins. Biochim. Biophys. Acta - Biomembr. 2015, 1848, 2839–2848.

60. Hoshi, T.; Zagotta, W.N.; Aldrich, R.W. Biophysical and molecular mechanisms of Shaker potassium channel inactivation. Science 1990, 250, 533–8.

61. Magidovich, E.; Orr, I.; Fass, D.; Abdu, U.; Yifrach, O. Intrinsic disorder in the C-terminal domain of the Shaker voltage-activated K+ channel modulates its interaction with scaffold proteins. Proc. Natl. Acad. Sci. U. S. A. 2007, 104, 13022–7.

62. Dosztányi, Z.; Mészáros, B.; Simon, I. ANCHOR: web server for predicting protein binding regions in disordered proteins. Bioinformatics 2009, 25, 2745–2746.

63. Sugase, K.; Dyson, H.J.; Wright, P.E. Mechanism of coupled folding and binding of an intrinsically disordered protein. Nature 2007, 447, 1021–1025.

64. Gasparini, A.; Tosatto, S.C.E.; Murgia, A.; Leonardi, E. Dynamic scaffolds for neuronal signaling: in silico analysis of the TANC protein family. Sci. Rep. 2017, 7, 6829.

65. Myrum, C.; Baumann, A.; Bustad, H.J.; Flydal, M.I.; Mariaule, V.; Alvira, S.; Cuéllar, J.; Haavik, J.; Soulé, J.; Valpuesta, J.M.; et al. Arc is a flexible modular protein capable of reversible self-oligomerization. Biochem. J. 2015, 468, 145–158.

66. Pastuzyn, E.D.; Day, C.E.; Kearns, R.B.; Kyrke-Smith, M.; Taibi, A. V; McCormick, J.; Yoder, N.; Belnap, D.M.; Erlendsson, S.; Morado, D.R.; et al. The Neuronal Gene Arc Encodes a Repurposed Retrotransposon Gag Protein that Mediates Intercellular RNA Transfer. Cell 2018, 172, 275–288.e18.

67. Gáspári, Z.; Nyitray, L. Coiled coils as possible models of protein structure evolution. Biomol. Concepts 2011, 2, 199–210.

68. Hayashi, M.K.; Ames, H.M.; Hayashi, Y. Tetrameric Hub Structure of Postsynaptic Scaffolding Protein Homer. J. Neurosci. 2006, 26, 8492–8501.

69. Hayashi, M.K.; Tang, C.; Verpelli, C.; Narayanan, R.; Stearns, M.H.; Xu, R.-M.; Li, H.; Sala, C.; Hayashi, Y. The postsynaptic density proteins Homer and Shank form a polymeric network structure. Cell 2009, 137, 159–71.

70. Wu, Q.; Sun, M.; Bernard, L.P.; Zhang, H. Postsynaptic density 95 (PSD-95) serine 561 phosphorylation regulates a conformational switch and bidirectional dendritic spine structural plasticity. J. Biol. Chem. 2017, 292, 16150–16160.

71. Serber, Z.; Ferrell, J.E. Tuning Bulk Electrostatics to Regulate Protein Function. Cell 2007, 128, 441–444.

72. Sun, Q.; Jackson, R.A.; Ng, C.; Guy, G.R.; Sivaraman, J. Additional Serine/Threonine Phosphorylation Reduces Binding Affinity but Preserves Interface Topography of Substrate Proteins to the c-Cbl TKB Domain. PLoS One 2010, 5, e12819.

73. Coba, M.P. Regulatory mechanisms in postsynaptic phosphorylation networks. Curr. Opin. Struct. Biol. 2019, 54, 86–94.

74. Davey, N.E.; Cyert, M.S.; Moses, A.M. Short linear motifs - ex nihilo evolution of protein regulation. Cell Commun. Signal. 2015, 13, 43.

75. Zeng, M.; Shang, Y.; Araki, Y.; Guo, T.; Huganir, R.L.; Zhang, M. Phase Transition in Postsynaptic Densities Underlies Formation of Synaptic Complexes and Synaptic Plasticity. Cell 2016, 166, 1163–1175.e12.

