## Supplementary Material for "Occurrence of ordered and disordered structural elements in postsynaptic proteins supports optimization for interaction diversity"

### Supplementary Discussion

#### *Protein-protein Interactions*

PPIs can also occur between ordered segments. The amyloid beta precursor protein is a receptor helping neurite growth and neuronal adhesion. The structure of the E2 domain of the protein was solved multiple times, both in dimeric [1] and monomeric [2] forms, suggesting the domain is stable in both forms. It was proposed that dimerization aids the masking of several sites with biological activity [3], and thus the oligomerization state may regulate fibroblast growth by hiding a short sequence motif (RERMS), which was shown to promote growth processes [4].

A particular case of domain-domain interaction is the complex formation by transmembrane (TM) proteins. Membrane proteins often assemble into higher-order complexes to mediate transport or signaling. Nicotinic acetylcholine receptors are ion channels assisting fast chemical neurotransmission with high functional diversity, assembling into a pentameric form. Monomers consist of four TM helices, and the complex has an inner ring of helices, shaping the pore, while the outer three helices of each monomer shield the inner rings from lipids. In the closed state, the inner helices are in proximity and gate the channel. Upon acetylcholine binding, a conformational change occurs: the individual TM subunits remain rigid, however, the arrangement of the channel changes to become permeable for sodium ions [5].

An interesting, however somewhat logical aspect of the TM proteome of PSD is the underrepresentation of different GPCR proteins, otherwise constituting the most abundant protein family [6]. Over half of GPCRs encode olfactory receptors, responsible for perceiving a high variety of chemicals. Cell-cell communication relies on receptors, however, the highly specialized ways of synaptic transmission may not require to distinguish such a diverse range of ligands as olfactory perception.

A more direct way to modulate PPIs is the control of the presence of its components at specific locations. Although it is hard to asses large scale protein synthesis and expression level data, ubiquitination sites hint the possibility of proteasome activity and degradation. It was shown that blocking either the synthesis or the degradation of synaptic proteins impairs the maintenance of late-phase long-term potentiation (L-LTP), however simultaneous blockade of protein degradation and protein translation leads to the rescue of L-LTP [7], suggesting an important role of ubiquitination in the PSD.

**Supplementary Figures**

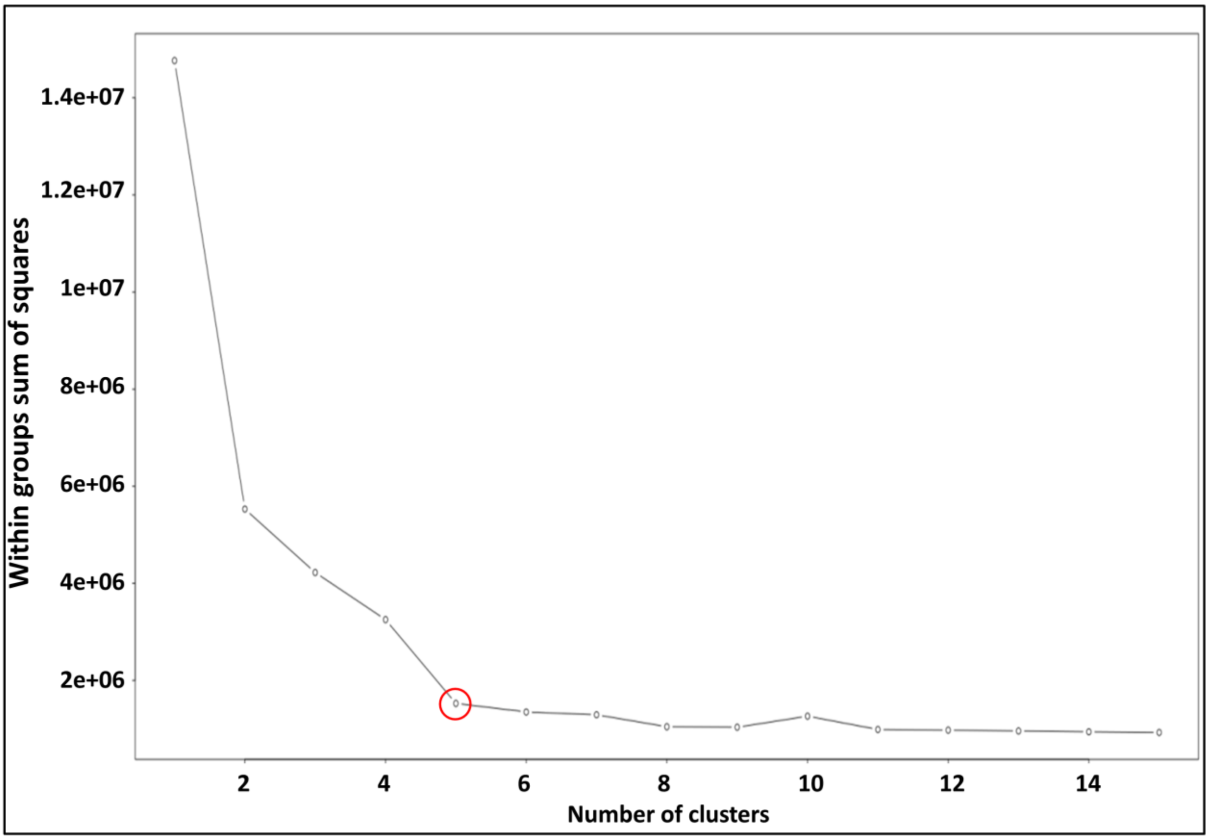


*SFigure 1: K-means evaluation of various numbers of clusters of AAIndex.*

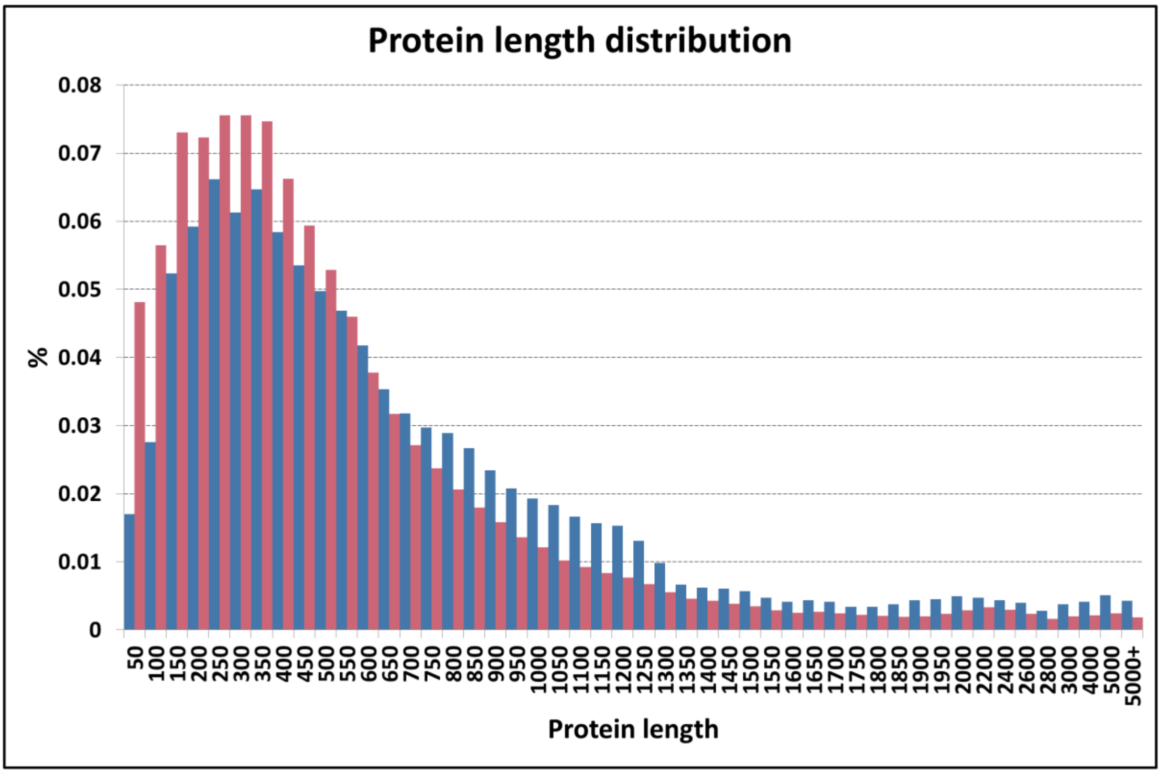


*SFigure 2: Protein length distribution (blue: PSD, red: human proteome)*

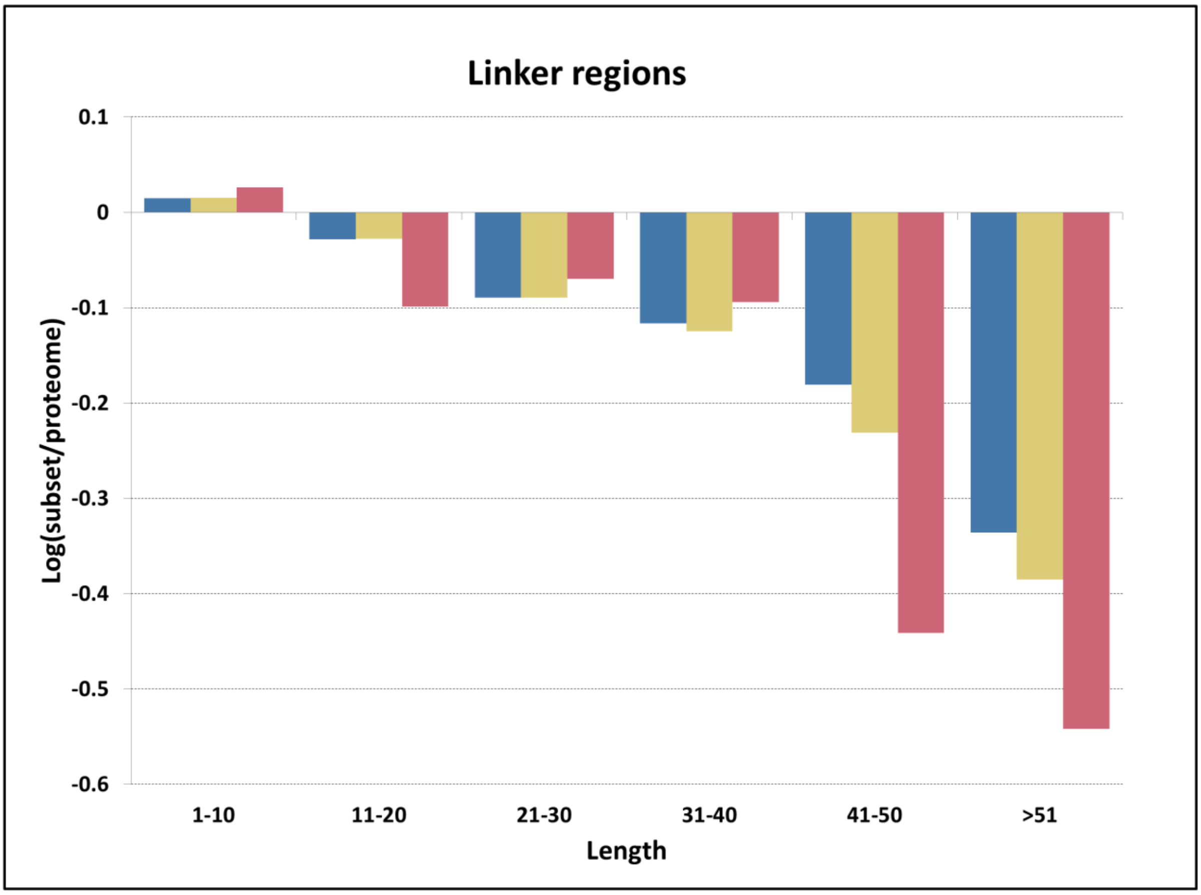


*SFigure 3: Flexible linker length distribution compared to the human proteome (blue: synaptome, yellow: PSD, red: PSC)*

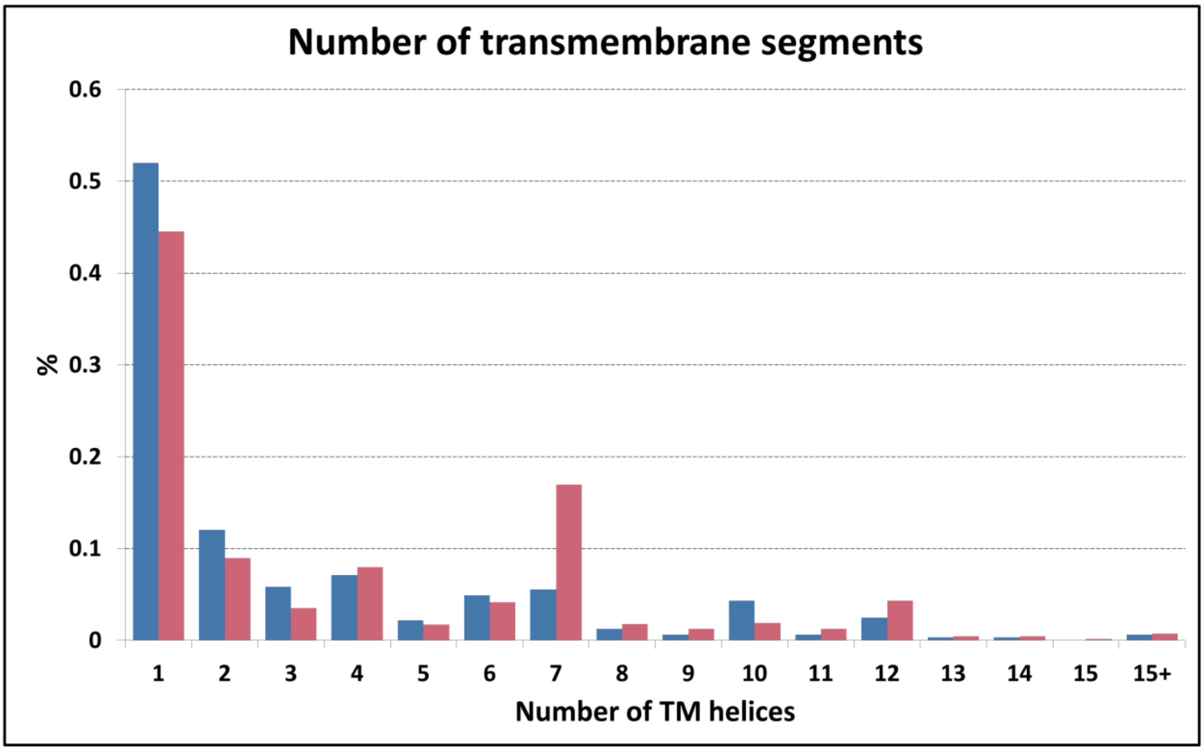


*SFigure 4: Transmembrane helix distribution (blue: PSD, red: human proteome)*

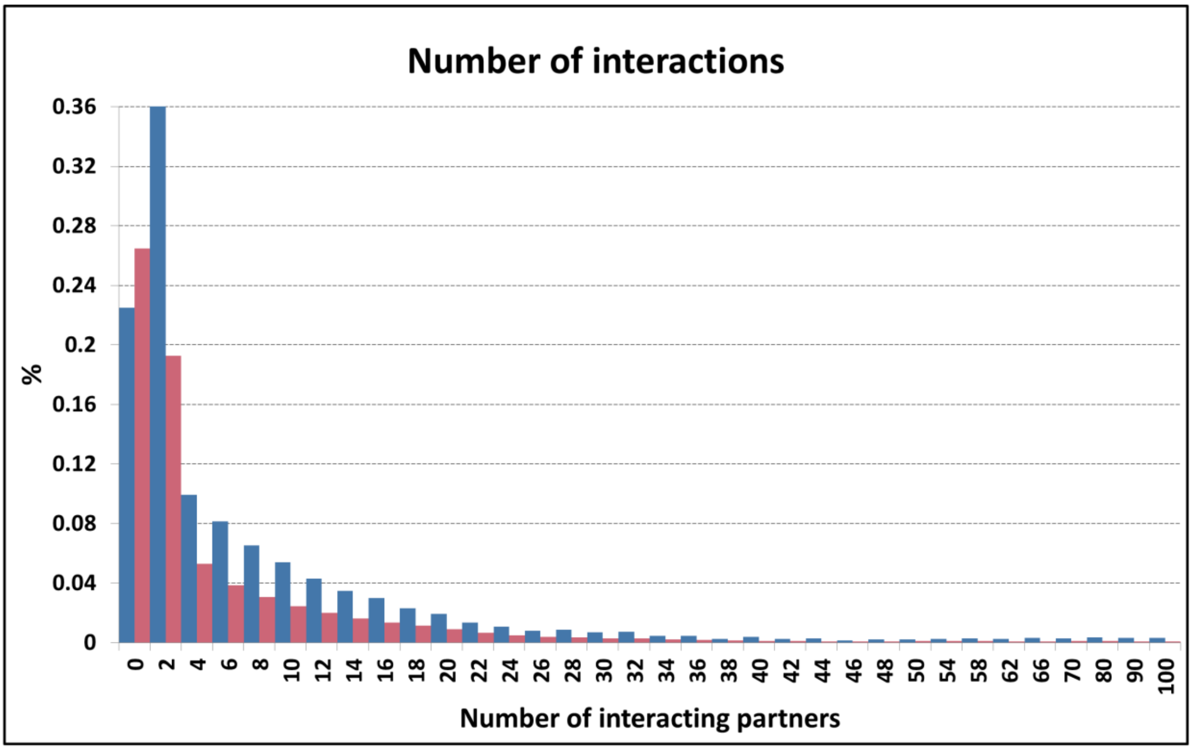


*SFigure 5: Distribution of the number of interacting partners (blue: PSD, red: human proteome)*

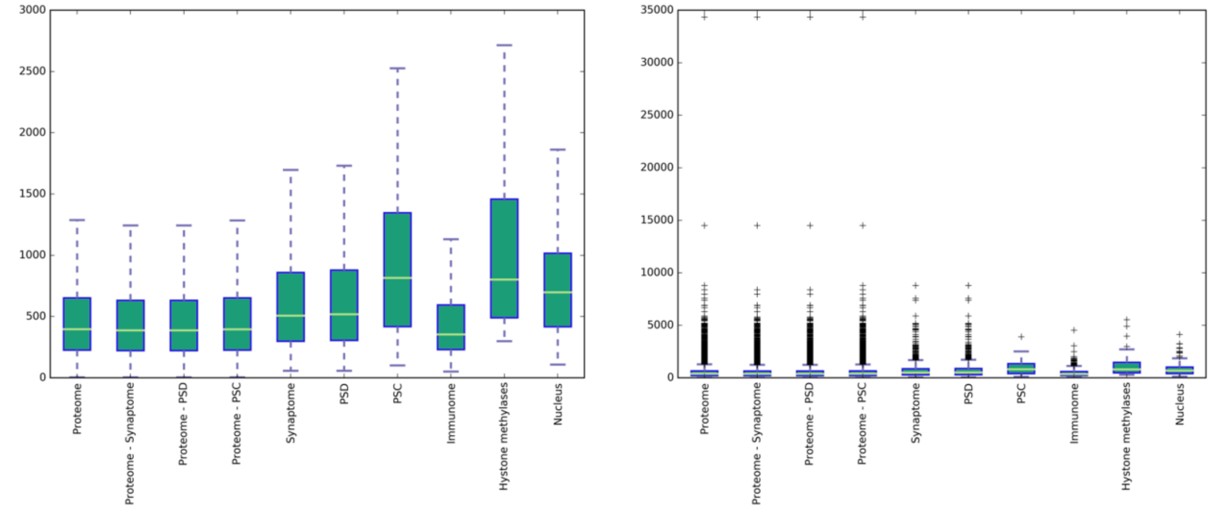


*SFigure 6: Boxplot of protein lengths in all datasets (left: without data points; right: with outlier data points. Note the different y-axes).*

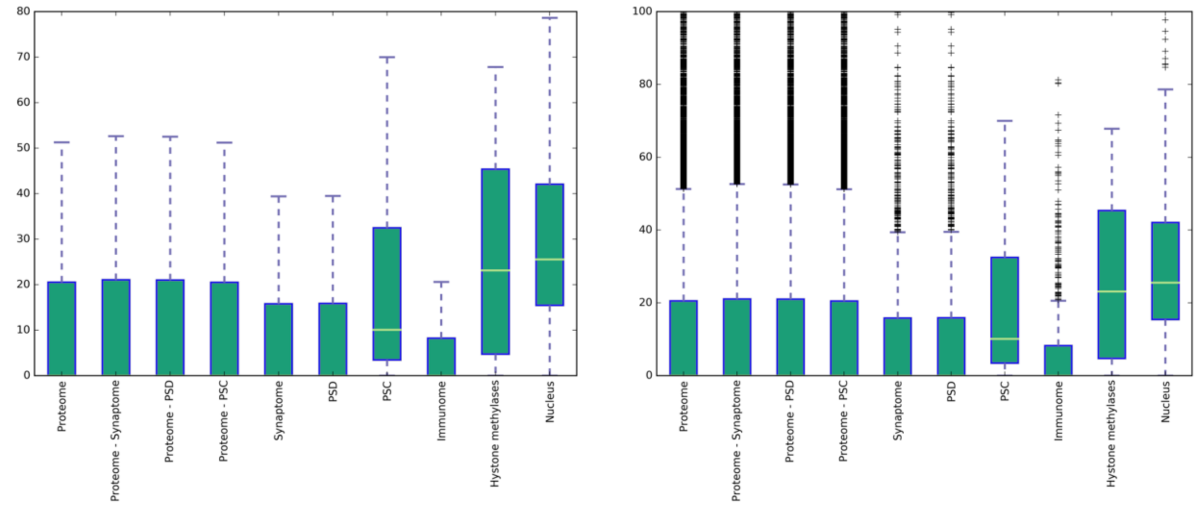


*SFigure 7: Boxplot of intrinsically disordered residue content in all datasets (left: without data points; right: with outlier data points. Note the different y-axes).*

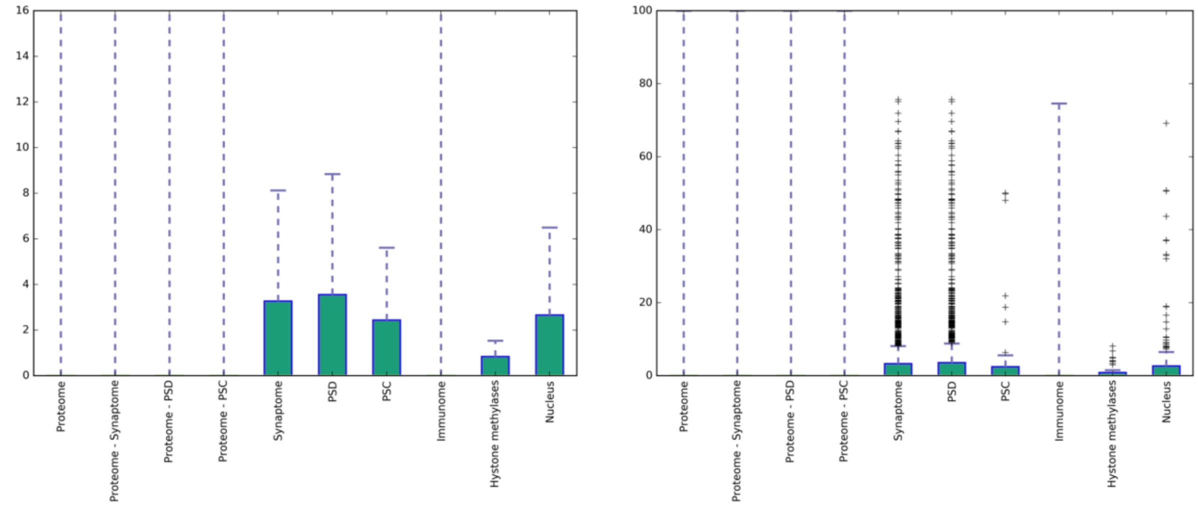


*SFigure 8: Boxplot of coiled-coil residue content in all datasets (left: without data points; right: with outlier data points. Note the different y-axes).*

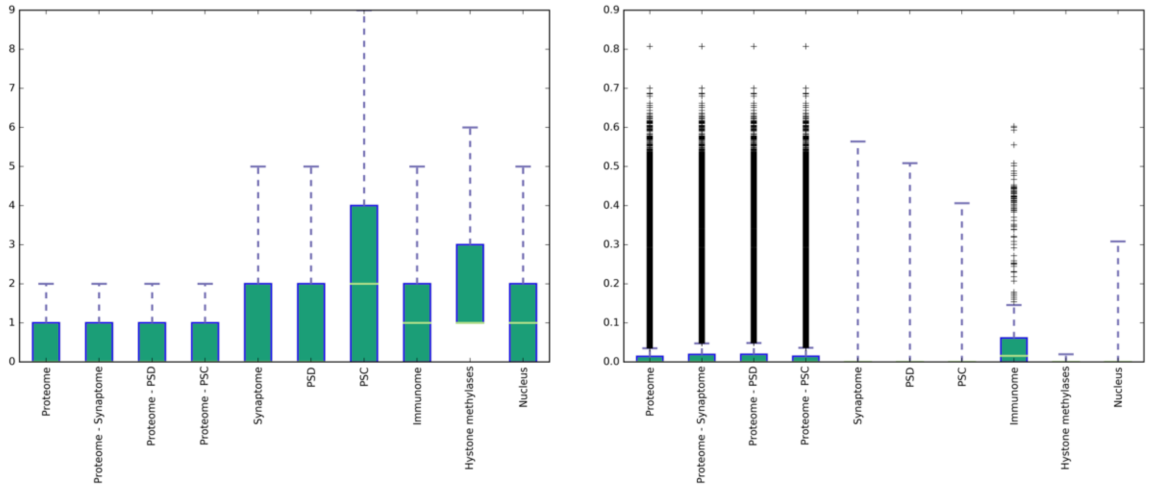


SFigure 9: Boxplot of transmembrane residue content in all datasets (left: without data points; right: with outlier data points. Note the different y-axes).

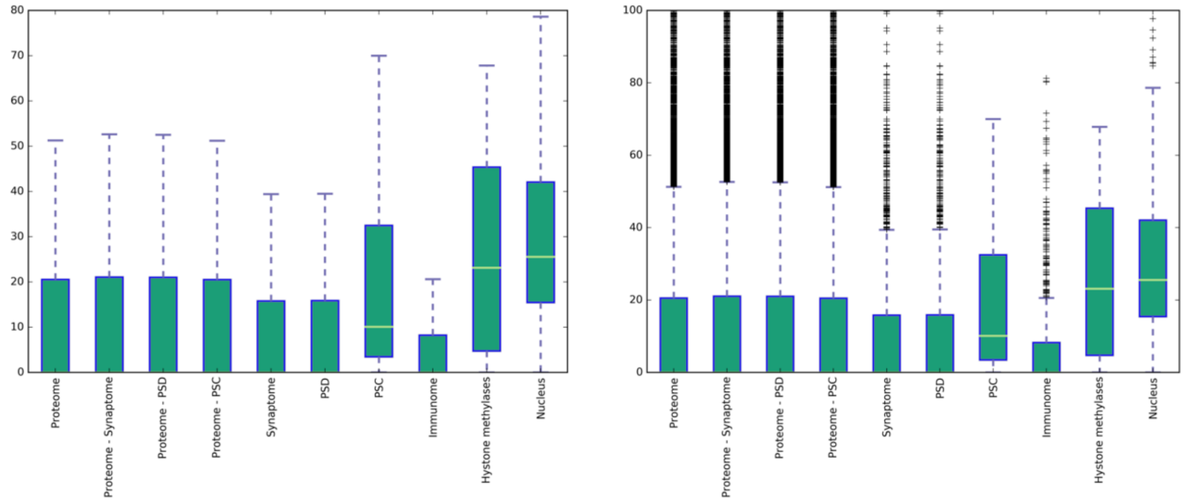


SFigure 10: Boxplot of the number of globular domains in all datasets (left: without data points; right: with outlier data points. Note the different y-axes).

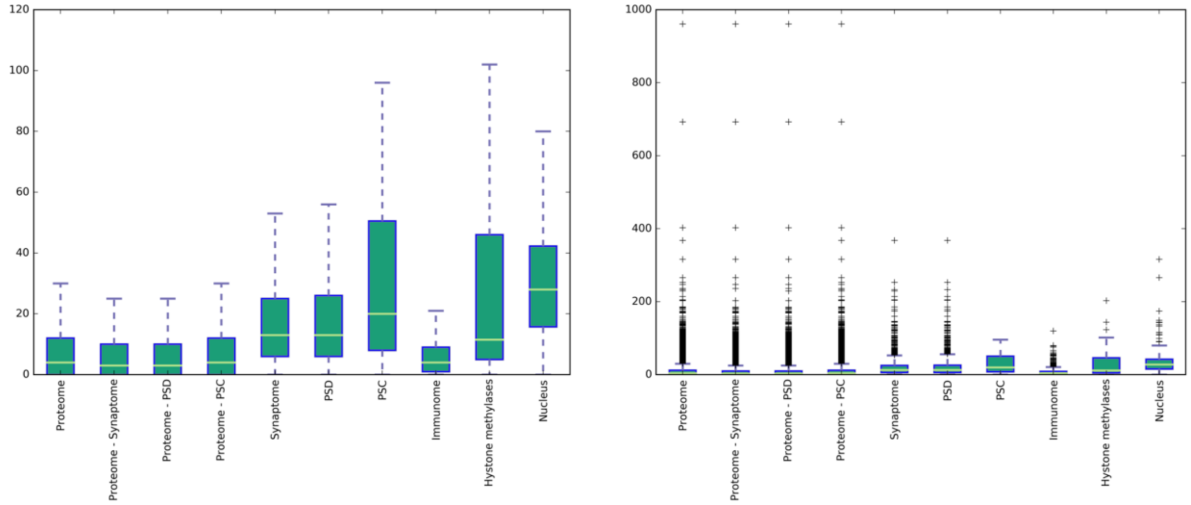


*SFigure 11: Boxplot of the number of phosphorylation sites in all datasets (left: without data points; right: with outlier data points. Note the different y-axes).*

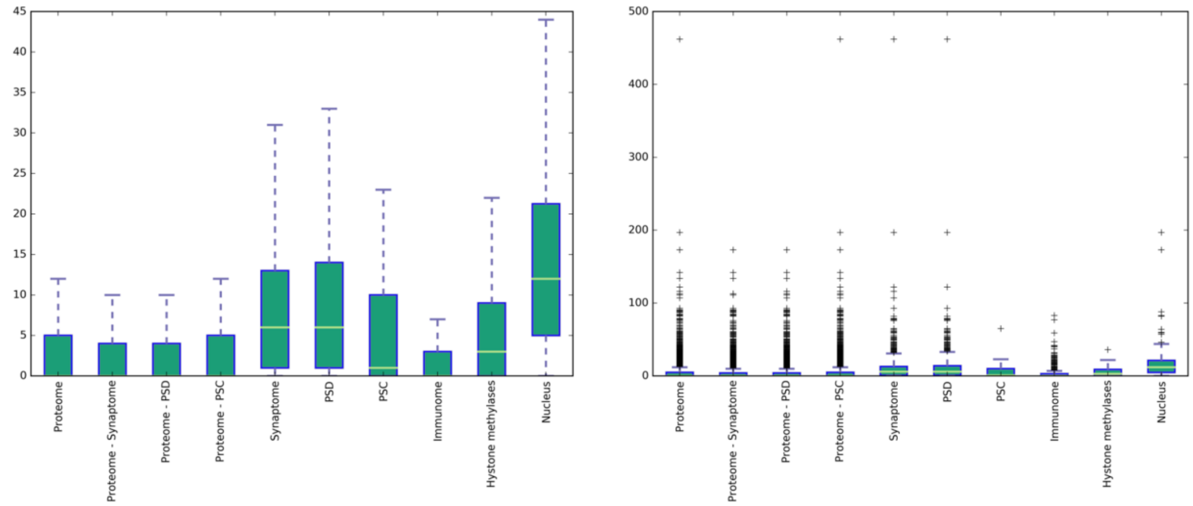


*SFigure 12: Boxplot of the number of ubiquitination sites in all datasets (left: without data points; right: with outlier data points. Note the different y-axes).*

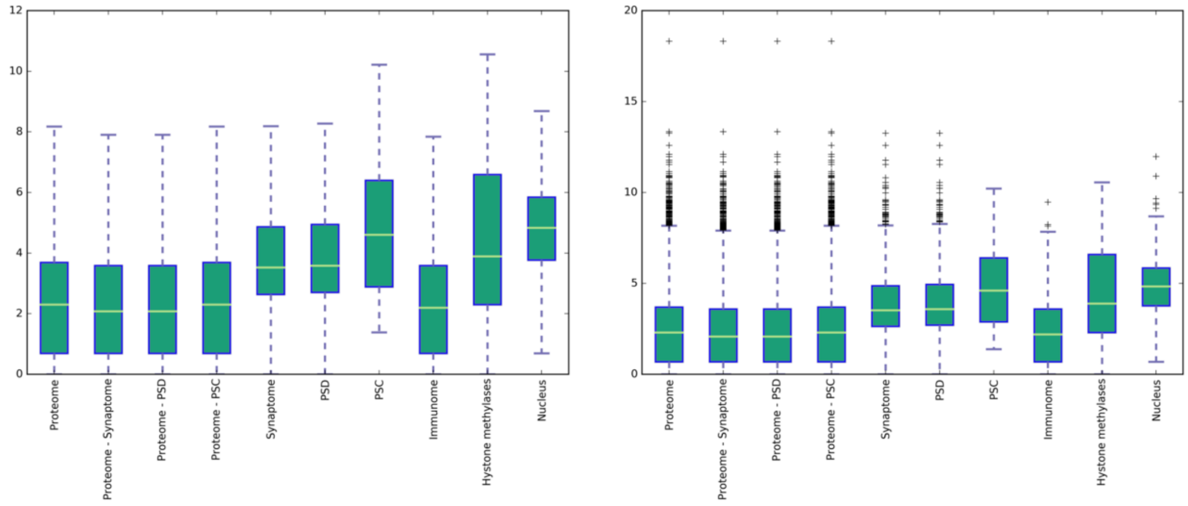


*SFigure 13: Boxplot of DPI values in all datasets (left: without data points; right: with outlier data points. Note the different y-axes).*

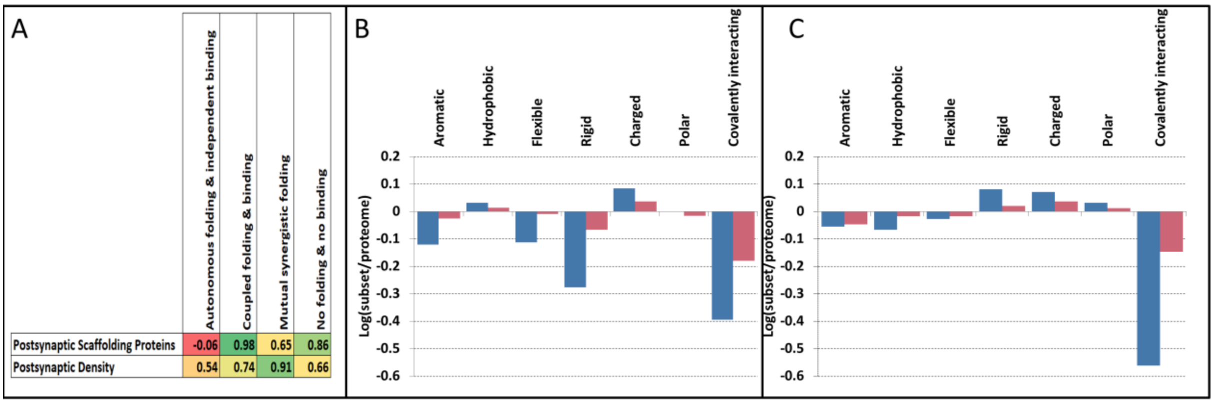


*SFigure 14:* A: Correlation of amino acid content between interaction classes based on the structural state of participating partners and postsynaptic proteins. Columns: Autonomous folding and independent binding (i.e., the binding of two or more ordered proteins), coupled folding and binding (where an ordered protein stabilize an IDP partner) and mutual synergistic folding (interactions formed exclusively by disordered proteins), No folding, no binding (i.e., the “classical” disordered definition). Rows: Postsynaptic Scaffold proteins, and proteins from the postsynaptic density. Color scales from red (negative correlation) to green (positive correlation). B: Change of amino acid content of the group mutual synergistic folding (blue) and PSD proteins (red) compared to the proteome. C: Change of amino acid content of the group coupled binding and folding (blue) and PSC proteins (red) compared to the proteome.

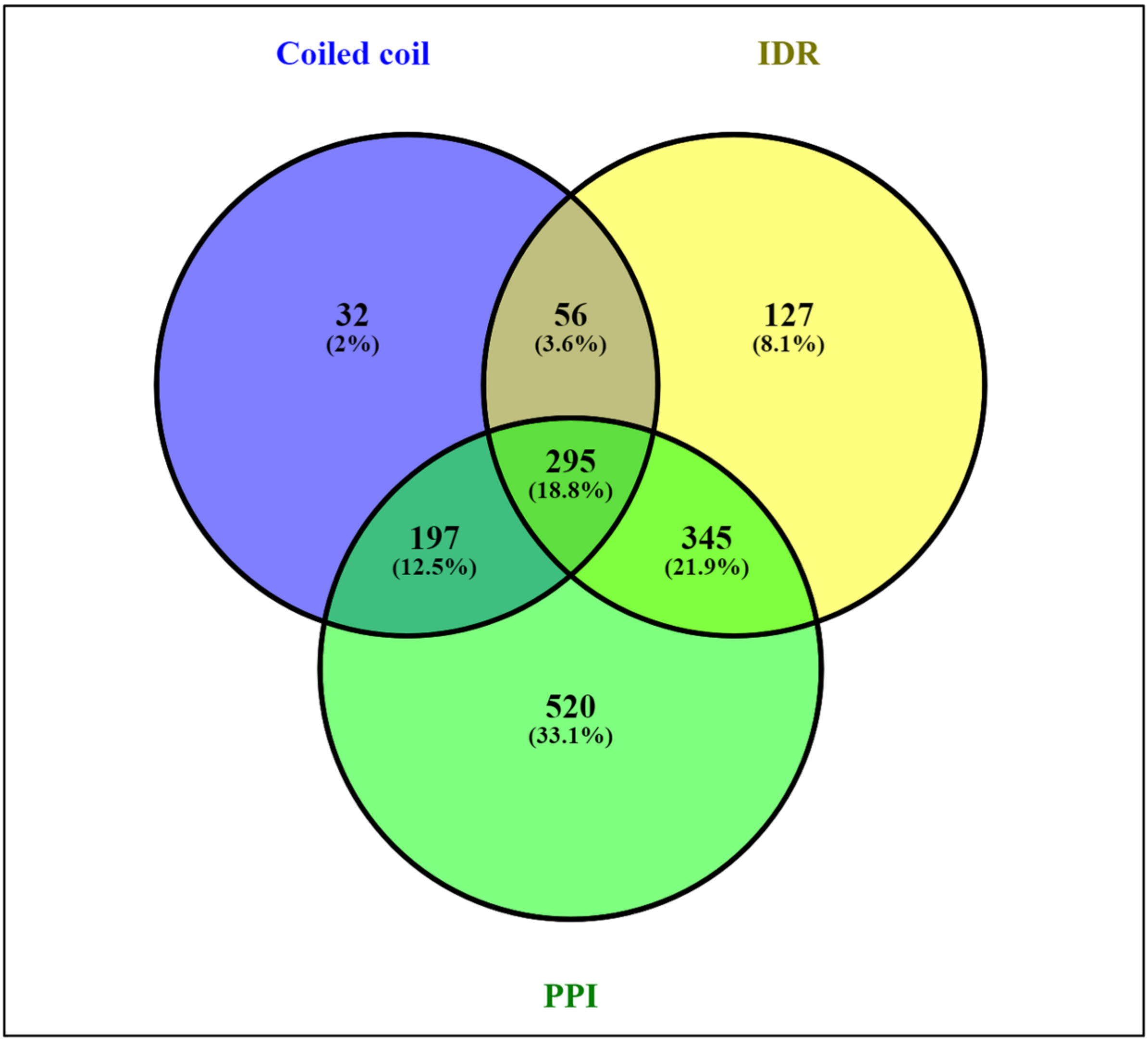


*SFigure 15: Overlap of coiled coils, IDRs and PPIs in PSD proteins.*

**Supplementary References**


1. Ninomiya, H.; Roch, J.M.; Sundsmo, M.P.; Otero, D.A.; Saitoh, T. Amino acid sequence RERMS represents the active domain of amyloid beta/A4 protein precursor that promotes fibroblast growth. *J. Cell Biol.* **1993**, *121*, 879–86.

2. Hoopes, J.T.; Liu, X.; Xu, X.; Demeler, B.; Folta-Stogniew, E.; Li, C.; Ha, Y. Structural Characterization of the E2 Domain of APL-1, a Caenorhabditis elegans Homolog of Human Amyloid Precursor Protein, and Its Heparin Binding Site. *J. Biol. Chem.* **2010**, *285*, 2165–2173.

3. Lee, S.; Xue, Y.; Hu, J.; Wang, Y.; Liu, X.; Demeler, B.; Ha, Y. The E2 Domains of APP and APLP1 Share a Conserved Mode of Dimerization. *Biochemistry* **2011**, *50*, 5453–5464.

4. Ninomiya, H.; Roch, J.M.; Sundsmo, M.P.; Otero, D.A.; Saitoh, T. Amino acid sequence RERMS represents the active domain of amyloid beta/A4 protein precursor that promotes fibroblast growth. *J. Cell Biol.* **1993**, *121*, 879–86.

5. Morales-Perez, C.L.; Noviello, C.M.; Hibbs, R.E. X-ray structure of the human α4β2 nicotinic receptor. *Nature* **2016**, *538*, 411–415.

6. Dobson, L.; Reményi, I.; Tusnády, G.E. The human transmembrane proteome. *Biol. Direct* **2015**, *10*, 31.

7. Fonseca, R.; Vabulas, R.M.; Hartl, F.U.; Bonhoeffer, T.; Nägerl, U.V. A Balance of Protein Synthesis and Proteasome-Dependent Degradation Determines the Maintenance of LTP. *Neuron* **2006**, *52*, 239–245.
